# Molecular insights into inhibitor action on a key bacterial metabolic enzyme *Cystathionine β-Synthase*

**DOI:** 10.1101/2025.07.03.662948

**Authors:** Sainath Polepalli, Anupam Roy, Bapan Mondal, Amit Singh, Somnath Dutta

**Author notes:** Corresponding authors. (SD); (AS). Contributed equally (Sainath Polepalli, Anupam Roy).

## Abstract

Tuberculosis (TB) remains a major global health threat, with Mycobacterium tuberculosis (*Mtb*) infecting nearly a quarter of the global population. Drug-resistant TB and HIV-TB co-infections emphasize the need for novel therapeutic approaches targeting essential metabolic pathways. Here, we investigated *Mtb* cystathionine β-synthase (*Mtb*CBS), a PLP-dependent enzyme critical for sulfur metabolism and redox regulation, owing its potential as a therapeutic target. We present the first high-resolution cryo-EM structure of *Mtb*CBS bound to aminooxyacetic acid (AOAA), and employed molecular dynamics (MD) simulations, quantum mechanics/molecular mechanics (QM/MM) calculations, and comparative inhibition studies to reveal the molecular basis and determinants governing irreversible PLP-enzyme inhibition. Our Cryo-EM structural analysis revealed two highly conserved active-site residues, T75 and Q147, critically stabilizing the inhibitor complex. Through molecular mimic studies, we demonstrated the precise structural factors and electronic features critical for inhibition efficiency. These findings provide the first mechanistic rationale for PLP enzyme inhibition and offer a generalizable framework for designing covalent inhibitors targeting PLP-dependent enzymes implicated in infectious diseases, cancer, and neurological disorders.

Graphical Abstract.
Proposed mechanism of *Mtb*CBS inhibition by AOAA. Proposed mechanism of action of AOAA illustrated through a graphical representation. The cartoon shows the domains of *Mtb*CBS. Tetrameric structure of *Mtb*CBS with active site highlighted (in red dashes). Enlarged view indicating the PLP-AOAA blocked intermediate interacting with residue Q147. Reaction scheme depicting the previous missing understanding of active site residue role in β-elimination. Schematic representation of AOAA inhibition mechanism in the catalytic cycle of *Mtb*CBS. Reactions were drawn using ChemDraw. The proposed scheme highlights that *AOAA* (highlighted Red) binding is stabilized by nearby residues Q147 and T75, leading to the formation of PLP-AOAA covalent adduct, and substrates (highlighted green) were incapable of reverting the suicidal inhibition. The directionality and reversibility were mentioned for each step.

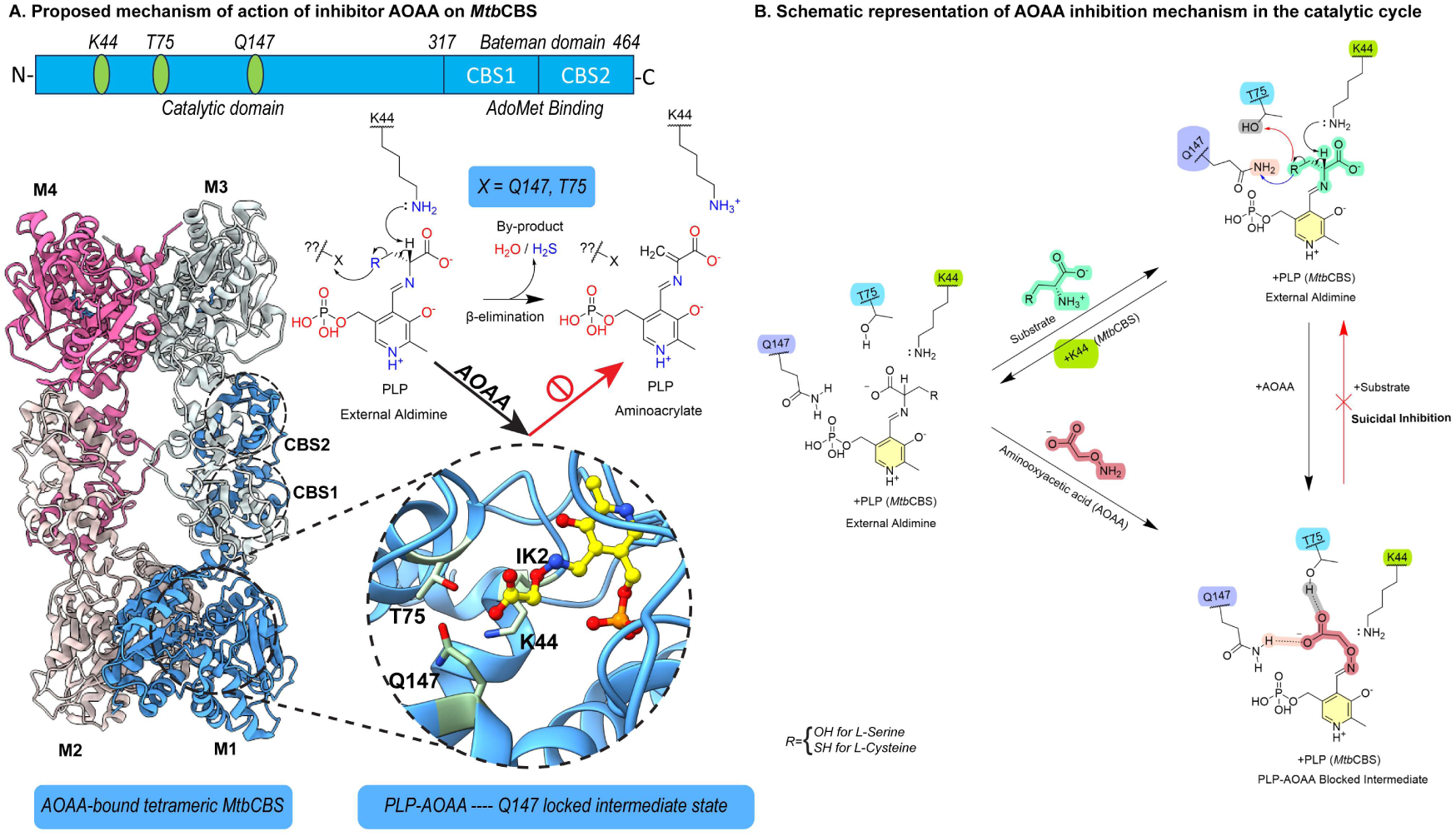

## Introduction

Antibiotic resistance remains a growing global threat due to emerging antimicrobial resistance (AMR) bacteria towards conventional antibiotics ^1^. In *Mycobacterium tuberculosis* (*Mtb*), available first-line drugs Isoniazid, Rifampicin, Ethambutol, and Pyrazinamide target enzymes that are involved in mycolic acid biosynthesis, cell wall synthesis, transcription, translation and energy production, respectively ^2,3^. However, *Mtb* has acquired resistance to those drugs (MDR-TB) and other second-line injectables (XDR-TB), making it a severe issue for the global population ^4^. Therefore, exploring more novel drug candidates targeting several other critical metabolic pathways is equally essential. The reverse trans-sulfuration pathway (RTS) is a metabolic process that assists *Mtb* in combating oxidative stress and survival during HIV-TB coinfection ^5^. Cystathionine β-Synthase (CBS) is the first enzyme in the RTS pathway responsible for the metabolism of sulfur-containing amino acids in a pyridoxal 5’-Phosphate- (PLP) dependent manner ^6,7^. Briefly, homocysteine generated from the methionine cycle is processed into L-cystathionine (Cysth) by chemical condensation with L-serine/L-cysteine (L-Ser/L-Cys) **(Fig 1A)** generating water/hydrogen sulphide (H2S) as by-products ^8^. Cystathionine γ-lyase (CGL), another PLP-dependent metabolic enzyme, further breaks the Cysth into L-Cys and α-ketobutyrate ^9^. While in an alternative de-sulfuration pathway, CBS and CGL convert L-Cys to L-lanthionine (Lanth) or L-Ser, leaving H2S in the cellular milieu ^10^. Thus, the RTS pathway significantly contributes to cellular redox homeostasis, thus facilitating the *Mtb* pathogenesis ^11,12^. Hence, *Mtb* Cystathionine β-Synthase (*Mtb*CBS) and *Mtb* Cystathionine γ- lyase (*Mtb*CGL) could be the novel drug targets for developing therapeutics against *Mtb*. Generally, designing new therapeutic agents leverages the knowledge of under-explored molecules and re-engineering their chemical architecture ^13^. However, understanding the mode of action of several such under-explored drug candidates is often hindered by limited structural information and structure-function cross-correlations ^14^. One such drug candidate, O-carboxymethyl hydroxylamine (AOAA, U-7524), used in clinical trials for Huntington’s disease, Colon cancer, and Tinnitus disease, was also reported to inhibit CBS enzymes ^15–17^. Molecular understanding of AOAA as a CBS inhibitor has been limited to cellular and biochemical studies ^18,19^. CBS enzymes were reported to catalyse a proposed β-elimination reaction in which the cofactor PLP formed two intermediates (aldimine and amino acrylate) species ^7,20^. Additionally, the diverse catalysis processes and versatile substrate preferences make targeting of PLP-based enzymes (class I-VII) challenging ^21^. Moreover, a detailed structural understanding, specifically regarding the role of active catalytic residues involved in the enzymatic process and their stereochemical preferences inside the active core, remains unexplored. In this context, we have recently resolved the full-length tetrameric structure of *Mtb*CBS along with the cryo-EM snapshots showing internal and external aldimine ^22^. In the present study, we structurally revealed a few functional side-chain residues at the active pocket that primarily accelerated the β-elimination process in *Mtb*CBS. We systematically deciphered the molecular chemistry behind the potency and efficacy of AOAA towards *Mtb*CBS enzyme. This was achieved by exploiting and decoding AOAA through Cryo-EM combined with structure-guided mutagenesis, in-vitro enzymatic assays, and in-silico investigations. Overall, this current investigation also provides significant molecular insights for translating new therapeutic candidates for other CBS enzymes

**Figure 1.**
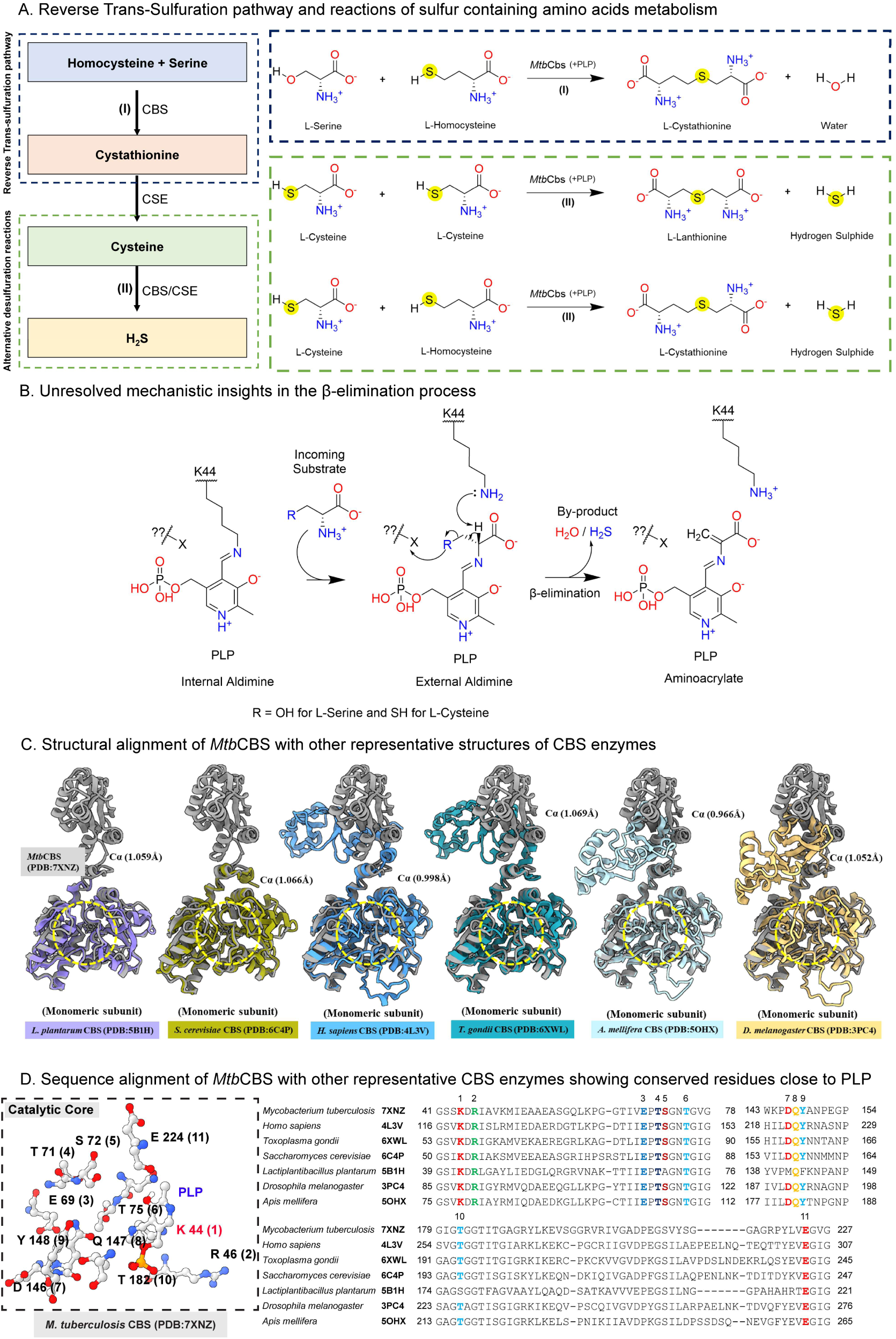
*Reverse Trans-sulfuration pathway steps, reactions, gaps and Conservation of enzymes*. ***A.* Reverse trans-sulfuration pathway steps and reactions involved in the metabolism of sulfur-containing amino acids.** A very simplified depiction of the sequential steps involved in the reverse trans-sulfuration pathway catalyzed by the CBS enzyme and in an alternative desulfuration pathway involving CBS/CSE enzymes to generate Hydrogen sulfide (H2S) is represented in the left panel. The right panel shows the trans-sulfuration reaction catalyzed by MtbCBS utilizing L-Serine and L-Homocysteine to generate L-Cystathionine (Marked in the blue box). The desulfuration reaction catalyzed by MtbCBS is marked in a green box showing the condensation of a molecule of L-Cysteine with another molecule of L-Cysteine or L-Homocysteine to produce L-Lanthionine or L-Cystathionine along with the by-product H2S. ***B.* Unresolved mechanistic insights in the β-elimination process.** Reaction schematic showing the β-elimination reaction in which the stabilizing nearby active site residues (X) and their role are marked with a question mark (?). ***C.* Structural conservation of active site architecture across CBS homologs.** Structural alignment of MtbCBS (7XNZ, Grey) with other representative structures of CBS enzymes from L. plantarum (5B1H, Light purple), S. cerevisiae (6C4P, Yellow green), H. sapiens (4L3V, Jordy Blue), T. gondii (6XWL, Picton Blue), A. mellifera (5OHX, Foam), D. melanogaster (3PC4, Golden brown). The calculated RMSD changes (in Å) (with respect to Cα) were written down in brackets in all cases. The conserved active site is marked by a yellow dashed circle. ***D.* Sequence conservation for the CBS enzymes from selective organisms.** The highly conserved polar & charged residues structurally vicinal to the PLP (-CHO) group (6Å distance) in the WT MtbCBS (PDB: 7XNZ) are shown in the left image and multiple sequence alignment in which the same residues highlighted separately using different colours. The PDB ID for the aligned proteins, along with the organism, were mentioned. Sequences were aligned using ClustalW. Residues were numbered on top and the corresponding numbers were provided in brackets next to the residues labelling.

## Results

Indeed, designing the novel targets specific to the CBS enzymes needs a robust structural overview that could explain the ongoing catalysis process. Trans-sulfuration pathways have been predicted to follow a series of chemical transformations. Therefore, a detailed understanding of the inhibitor actions through a correlated structure-functional relationship is required to boost inhibitor specificity and efficacy towards CBS. To prevail inhibitor/drug action, the detailed catalytic process and catalytic residues at the active enzymatic core are essential for this investigation.

### Catalytic cycle of CBS enzymes and missing details about functional side chains in the β- elimination process

CBS catalyzed cystathionine & lanthionine biosynthesis and de-sulfuration through a PLP- catalyzed β-elimination reaction **(Fig 1A)**. The holo-enzyme holds cofactor PLP in the active site via a covalent interaction (internal aldimine) with internal nucleophilic side chain lysine (Step I) **(Fig 1B & S1A)**. Our previous investigation and cryo-EM structural analysis found that K44 residues formed a stable internal aldimine adduct with aldehyde moieties of PLP in tetrameric *Mtb*CBS ^22^. The ζ- amine group of K44 residues acts as a competitor nucleophilic ligand/ base for any incoming monoamine substrate. However, the catalysis process proceeds in the presence of substrates homocysteine, cysteine and serine belonging to the monoamine class. The presence of substrate causes the chemical conversion from stable internal aldimine (PLP-Lys) to external aldimine (PLP-Substrate) ^6^, which proceeds via reaction intermediate gem-diamine formation (aldimination process, Step II) **(Fig S1A)**. Indeed, the β-elimination path is a crucial step that generates α-β unsaturated amino acrylate intermediate ^23^ (Step IV), primarily regulates the de-sulfuration process (H2S) and rate-determining for the cystathionine and lanthionine biosynthesis ^24^. The nucleophilic addition of another incoming substrate to this Cβ-position of PLP-aminoacrylate leads to the formation of PLP-Lnth/Cyth covalent adducts (Step V). Finally, the catalytic cycle reverts/restores after the elimination of product ^25^ caused by the nucleophilic substitution of internal lysine (trans-aldimination, Step VI). Lys residue close to PLP had been earlier suggested to participate in the reaction’s mechanism for *human* CBS, *L. plantarum* CBS and other PLP enzymes, respectively ^26,27^. From our previous investigation, we also observed that the K44A variant was incapable of reaction and unable to generate post-β-eliminated by-product H2S ^22^. Apart from external to internal aldimine conversion during product elimination from the active site, Lys- ζ NH2 was also predicted as a sensible nucleophile for withdrawing of proton (H^+^) from Cα of the external PLP-substrate aldimine complex (Step III), which initiated the β-elimination process. However, the removal or stabilization of -SH (cysteine)/-OH (serine) at the Cβ-position could also be accelerated by any other side chains of active catalytic residues **(Fig 1B)** that primarily regulate the enzymatic process remain unexplored for CBS enzymes. Furthermore, to get insights into the catalytic domain and the residues close to cofactor PLP, we performed structure-based-sequence alignment of available isoforms (crystal structure) from various organisms of CBS enzymes with the monomeric subunit of our previously resolved tetrameric Cryo-EM structure of holo- *Mtb*CBS (PDB ID: 7XNZ) **(Fig 1C)**. The structure-based sequence alignment revealed several sequentially conserved polar amino acids in the active core of the CBS enzymes. Herein, we depicted a few structurally conserved polar residues located close (within 6 Å) to coenzyme PLP in the active sites relative to *Mtb*CBS (K44, R46, S72, T75, D146, Q147, Y148, T183 and E224) **(Fig 1D, S1B)**. Except for nucleophilic lysine (universal base, K44 in *Mtb*CBS) residues close to cofactor PLP, the functional role of other conserved polar residues in the active pocket has not been known so far. Since an inhibitor for an enzyme could also perturb/impede the ongoing reaction steps, we hypothesize that exploiting inhibitor property in the *Mtb*CBS enzyme could elucidate both the catalytic role of active residues and unveil the inhibitor action in other CBS enzymes simultaneously. Therefore, to diagnose the impacts of inhibitors on PLP- catalyzed reaction in *Mtb*CBS, we have used a potential inhibitor, AOAA and conducted functional and cryo-EM-based structural studies. Initially, to characterize the impaired reaction step in *Mtb*CBS caused by AOAA, we performed several in-vitro enzymatic assays.

### AOAA restricted trans-aldimination process irreversibly

To gain insights into the reaction intermediates modulated by inhibitor AOAA, we purified and further functionally characterized tetrameric *Mtb*CBS as described in our lab-established workflow **(Fig S2A-B&D)** ^22^. The recombinant tetrameric *Mtb*CBS (200kDa) showed a substrate (L-Cys) concentration and SAM-dependent activity enhancement through monitoring the by-product (H2S) of the β-elimination reaction (Km & Vmax for +/- SAM) **(Fig S2C)**. The pre-β-eliminated intermediate (External aldimine) content was monitored through fluorescence measurements, while the post-β-eliminated by-product (H2S) was detected indirectly by lead acetate-based assay. Further, we treated freshly purified tetrameric *Mtb*CBS (10μM) with increasing concentrations of AOAA (20-200 μM) in reaction buffer, incubated for 10 minutes at 25°C and intensity in fluorescence emission corresponding to internal aldimine (λem=510nm) and oxime (λem=450nm) species were monitored spectroscopically. We observed a sequential reduction in aldimine peak intensity and a shift in peak towards oxime wavelength regime with increasing AOAA concentrations **(Fig S3A-B)**. The ability of AOAA to compete with the PLP- external aldimine intermediate was monitored in the presence of 10mM serine-treated *Mtb*CBS (10μM) with increasing AOAA concentrations (5-1000 μM), and further fluorescence emission was monitored. We observed a reduction in aldimine peak intensities and a concomitant increase in oxime peak intensity as a variant of AOAA concentration **(Fig S3D)**. Identical experimental trends were observed with another substrate, cysteine, where we monitored the external aldimine and post-β-eliminated by-product content **(Fig S3E)**. The IC50 of AOAA for our experimental study of serine-treated and cysteine-treated *Mtb*CBS was calculated as ∼100μM, respectively **(Fig S3F)**, suggesting AOAA inhibition is independent of the substrate’s nature/preferences. We further monitored the shift in fluorescence emission corresponding to the *Mtb*CBS (10μM) (pre-treated with 0-200μM AOAA) upon the addition of molar excess of L-Ser and L-Cys (10mM) separately **(Fig-2 A)**. However, we did not observe any further increase in aldimine species. No reduction in the observed oxime peak intensity suggested that suicidal inhibition was caused by AOAA treatment. This investigation indicated that AOAA could potentially (irreversibly) block cofactor PLP in Step I-II or Step III-IV. The mechanism behind AOAA’s suicidal inhibition, even in the presence of excess competitive monoamine substrates and a strong internal base (K44) that reversibly regulates the reaction, remained unclear. To determine whether the covalent binding of AOAA to PLP (Step I) alone is sufficient for suicidal inhibition or if additional interactions with active site side chains are necessary, we conducted a Cryo-EM structural analysis of AOAA-treated *Mtb*CBS **(Fig 2B-E)**.

**Figure 2.**
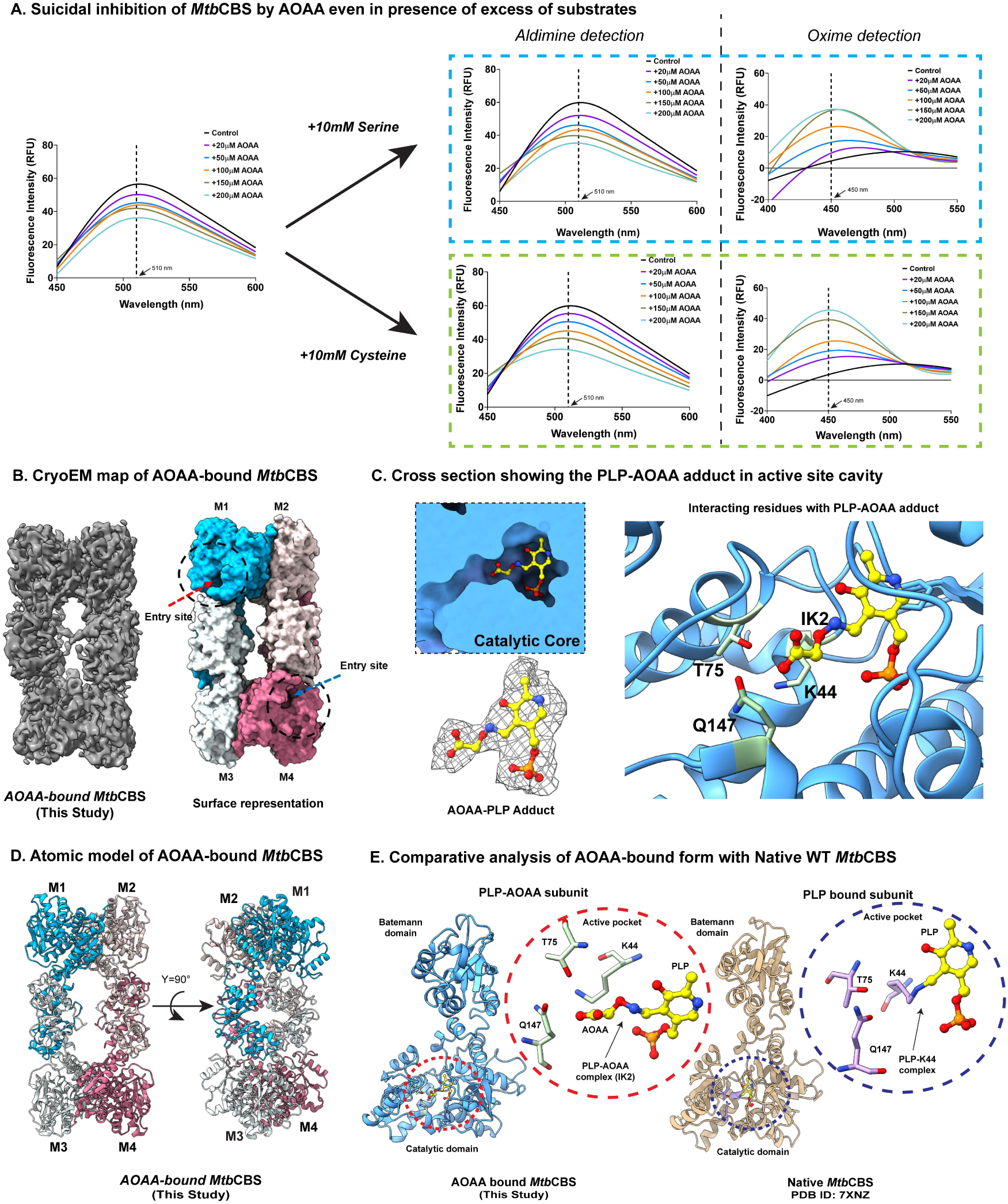
*Structural insights into inhibition mechanism by AOAA on MtbCBS* ***A.* Irreversible inhibition of tetrameric MtbCBS by AOAA.** Suicidal inhibition of MtbCBS by AOAA (See Methods) inferred by fluorescence emission spectra recorded from 450nm to 600nm upon exciting at 410nm of (Left panel) MtbCBS incubated with increasing concentration of AOAA (0-200μM) (Upper and lower panels). The ability of substrates to reverse back the MtbCBS activity was assessed by the addition of 10mM of Serine (Upper panel, blue dashed line) and 10mM of Cysteine (Lower panel, green dashed line) to the MtbCBS pre-treated with AOAA. The extent of changes in the aldimine and oxime species were inferred from the recorded fluorescence emissions. *B-E. Cryo-EM structural overviews to understand the effect of AOAA on MtbCBS***. *B.*** Cryo-EM density map of tetrameric MtbCBS along with the surface representation (coloured showing individual protomers) of the atomic model (indicating the active site in dashed lines) in the presence of CBS inhibitor AOAA. ***C.*** Cross section showing the PLP-AOAA adduct (IK2) in the active site, nearby interacting side chain residues in stick representation along with the AOAA-PLP cryo-EM density in mesh representation. ***D.*** Atomic model of the AOAA-bound MtbCBS structure (with individual protomers labelled). ***E.*** Comparative analysis of protomeric subunits of AOAA-bound (blue; PDB ID: 9U7O) with Native (Light brown; PDB:7XNZ) MtbCBS along with the active site arrangement (focusing T75, Q147 residues) in AOAA-bound (light green) and K44-PLP (magenta) MtbCBS structures.

### Cryo-EM structural insights of AOAA binding to WT-MtbCBS structure

To unravel more insights about AOAA interactions with PLP and nearby side chain residues in WT-*Mtb*CBS, especially if the enzyme undergoes a significant conformational change upon AOAA binding to the active site, we performed Cryo-EM investigation for the AOAA incubated sample WT-*Mtb*CBS (Methods). The preliminary NS-TEM analysis **(Fig S8C)** and cryo-EM reference-free 2D class averages **(Fig S4A)** predicted that the protein preserved its native tetrameric state in the presence of AOAA. From our 3D reconstituted cryo-EM density map (3.96 Å) and fitted atomic model **(Fig 2B&D)**, we further confirmed the intact homo- tetrameric state of the enzyme. We also have identified a continuous unmodelled (apart from CBS density) extra density connected directly with the free PLP ligand. We could successfully dock (fitted) the PLP-AOAA complex (4’-deoxy-4’-acetylyamino-pyridoxal-5’-phosphate, IK2) in the empty and unmodeled EM volume **(Fig 2C, Movie-1)**. We further compared the conformational changes in the PLP-AOAA complex in the core of *Mtb*CBS with our previously resolved holo- and serine-bound forms of *Mtb*CBS. Indeed, we did not observe significant differences in the individual protomeric structures of AOAA-bound (This study; PDB ID: 9U7O) and Native *Mtb*CBS (PDB ID: 7XNZ) **(Fig 2E)**. Closely examining active site PLP-Ser and PLP-AOAA adducts in their respective structures suggested that both maintain an identical chemical conformational space in the active site **(Fig 3A)**. However, PLP-AOAA would get additional stability due to the close encounter of terminal COO- of AOAA with polar residue Q147 (proximal contact) and T75, respectively. Therefore, we speculated that such additional interactions could further stabilise the PLP-AOAA complex strongly compared to PLP-substrate external aldimine adducts, which might explain the functional irreversibility of AOAA over substrates. Since residues T75 and Q147 were close to Cβ-OH of PLP-Serine adduct and highly conserved among CBS enzymes, we speculated that those residues might cause a weakening of Cβ-OH covalent bond and subsequent release of the functional moieties (-OH/-SH) from Cβ- position leaving reactive α-β unsaturated amino-acrylate. Therefore, we further anticipated that the PLP-AOAA complex could block the functional side chains facilitating the β-elimination process (Steps III &IV).

**Figure 3.**
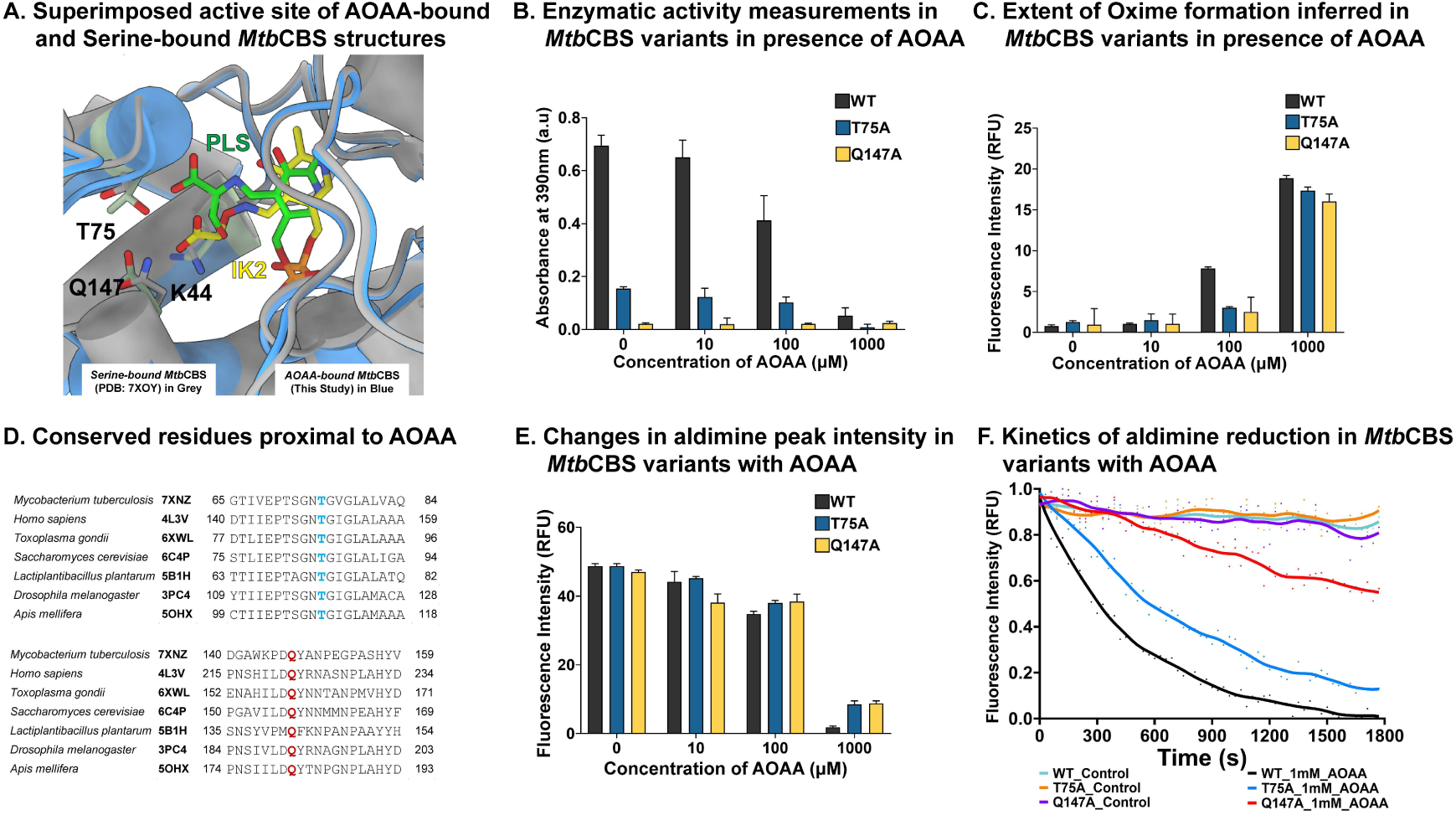
*Identifying functionally reactive residues by mutagenesis studies and its impact on AOAA binding*. ***A.*** Enlarged view of the conformational superposition of active sites of AOAA-bound (Blue; PDB 9U7O) and Serine-bound (Grey; PDB 7XOY) MtbCBS structures suggesting the close encounter of side chain T75 and Q147 with COO- a group of AOAA. Bar plots showing the changes in **B.** enzymatic activity, **C.** oxime formation in MtbCBS WT and mutant variants (T75A and Q147A) with different AOAA concentrations (10, 100, and 1000 μM). **D.** Sequence alignment depicting conservation in Thr-75 & Gln-147. **E.** Bar plot of aldimine changes with similar conditions in **B&C** (See Methods). **F.** Kinetics of aldimine reduction in WT and mutants with 1mM AOAA for a 30-minute interval.

### Identifying functional side chains for the β-elimination process and inferring their role in AOAA binding and function

To understand the role of these conserved residues in close vicinity to PLP-AOAA adduct **(Fig 3A&D)**, we individually generated T75A and Q147A mutants and assessed their role in the catalytic process. As these mutations are located in the core of the active site and not at the dimer or tetramer interfaces, we expect all mutant variants of *Mtb*CBS to maintain the native tetrameric structure, similar to the wild-type enzyme. The presence of tetramers was confirmed in our gel-filtration and NS-TEM study **(Fig S8A-B)**. We further observed that the T75A mutation reduced the enzymatic activity (PbS assay; λabs=390nm, cysteine as substrate) by 80% compared to wild-type (WT) *Mtb*CBS, while the alanine substitutions at the Q147 residue retained only 10% of the activity **(Fig 3B)**. Additionally, we conducted these experiments at varying AOAA concentrations (10, 100, 1000 μM) to assess its effect on the mutants. The results indicated that T75 and Q147 are critical for the β-elimination process and irrespective of AOAA. To determine whether T75 or Q147 residue is involved in AOAA binding, we monitored changes in oxime formation (λmax=450nm) and the relative reduction in external aldimine (λmax =510nm) (PLP-Serine; external aldimine). At lower AOAA concentrations (10μM), no significant differences in oximes formation were observed between the mutants and wild-type (WT) *Mtb*CBS **(Fig 3C)**. Similar trends were observed in aldimine intensities as well **(Fig 3E)**. At near IC50 concentrations (100μM), we noted a significant decrease in oxime formation and a partial increase in aldimine intensity in the mutants compared to WT *Mtb*CBS. In samples treated with 1mM AOAA, a substantial reduction in aldimine and increased oxime formation were observed in WT *Mtb*CBS relative to the mutants. The role of side chains in AOAA binding was further supported by kinetic results, which showed a slower rate of aldimine reduction in both mutants relative to WT *Mtb*CBS **(Fig 3F)**.

Based on our enzyme activity and structural investigations, we hypothesize that the relatively more polar residue Q147 plays a role in the reaction and AOAA binding. To test this, we mutated Q147 to less polar residues (Q147Y, Q147N, and Q147E) **(Fig 4A)**. While Q147N retained 10% of the activity, no detectable H2S production was observed for Q147Y within the same reaction time **(Fig 4B)**. However, a small amount of lead-sulphide formation (λabs=390nm) was detected for Q147N alone (∼10% of WT *Mtb*CBS) after approximately 3 hours **(Fig 4C)**. Aldimine rate kinetics data revealed reduced oxime formation in Q147N/E mutants, further supporting Q147’s involvement in AOAA binding. Notably, no significant reduction in the internal aldimine was observed for the Q147E (charge reversal) mutant in the presence of 1mM AOAA over the entire detection period **(Fig 4D)**. Therefore, if the terminal carboxylate group in AOAA was solely responsible for binding (Cryo-EM and mutagenesis studies) with the polar side chain of Q147 and T75, which stabilized the PLP-oxime adduct in the active site, we hypothesize that deleting the carboxylate group from AOAA would likely weaken both its binding and inhibition efficiency.

**Figure 4.**
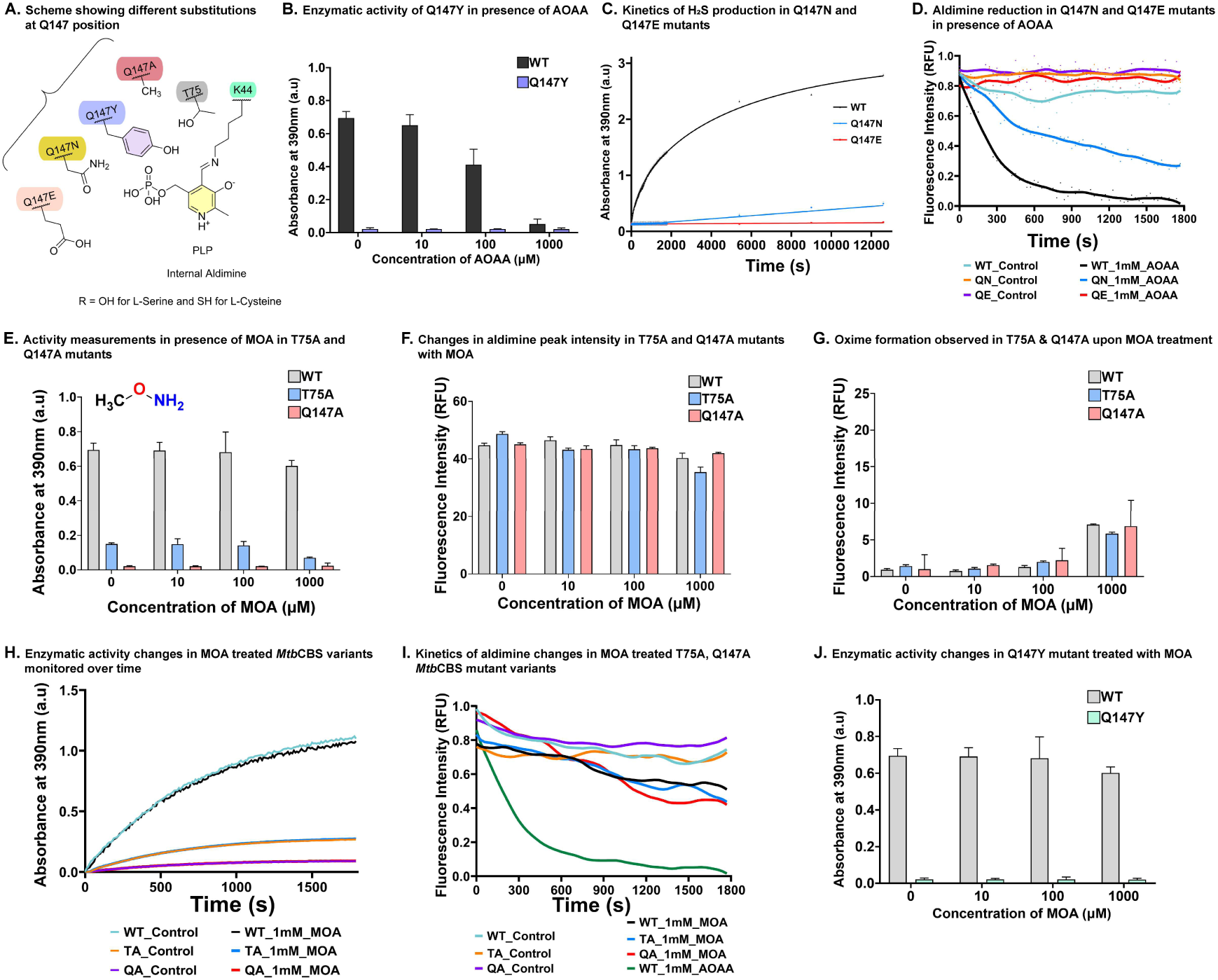
*Significance of Q147 residue in reaction and AOAA binding via terminal carboxylate group*. **A.** Scheme highlighting the different amino acid substitutions (A, Y, N, E) at Q147 position. **B.** Enzymatic activity of Q147Y mutant in the presence of AOAA measured through lead acetate-based assay. H2S production is below detectable levels in the Q147Y mutant. **C.** Kinetics of H2S production monitored in Q147N and Q147E mutants continuously for the first 30-minute period, followed by detection at every 1h interval. **D.** Aldimine changes in Q147N and Q147E mutants compared to wild type upon 1mM AOAA treatment. Mutants show a lack of effective reduction in aldimine during detection. Bar plots showing changes in **E.** Enzymatic activity, **F.** Aldimine peak intensity and **G.** extent of oxime formation in T75A and Q147A mutants with different concentrations (10, 100, and 1000 μM) of Methoxyamine (MOA) that mimic the absence of terminal carboxylate in AOAA. **H-I.** Kinetics of enzymatic changes and aldimine changes in T75A and Q147A mutants treated with 1mM MOA analogue. **J.** Bar plot indicating lack of observed enzymatic H2S production in Q147Y treated with different concentrations of MOA.

### Terminal Carboxylate group in AOAA is crucial for binding with active side chains and inhibition efficiency

To validate this, we used an AOAA-analogue molecule, Methoxyamine (MOA; an adjunct to alkylating agents in anti-cancer treatment ^28^), that lacks the terminal carboxylate group and conducted its functional activity relative to AOAA. Herein, we also maintained similar experimental conditions to those previously conducted for AOAA. In the case of MOA-treated WT *Mtb*CBS, we did not observe a significant reduction in the extent of aldimine and increased oxime formation even at 1mM concentrations compared to the AOAA-treated WT *Mtb*CBS sample **(Fig 4F-G)**. The lack of significant changes in pre-β-eliminated species indicated that AOAA became functionally inactive without its terminal carboxylate, as it could no longer bind to T75 and Q147 residues. Since the mutants were unable to generate H2S, no detectable H2S (PbS; λabs390nm) was observed, similar to AOAA-treated mutants **(Fig 4E&J)**. Similar results were inferred from the kinetic study for a period of 30 mins **(Fig 4H)**. These two residues (in WT *Mtb*CBS) are essential for AOAA binding but cannot interact with MOA (due to the deleted carboxylate), explaining the minimal reduction in aldimine even at high molar excess (1mM) MOA treatment in WT *Mtb*CBS. MOA binding was consistent across both WT *Mtb*CBS and mutant variants, as evidenced by the similar loss of aldimine content in kinetic studies **(Fig 4I)**. However, unlike AOAA, MOA cannot impede functional residues as substrates can reversibly displace MOA from the PLP-covalent linkage, restoring enzyme activity in the WT enzyme. These findings suggest that AOAA impacts both Step I and Step II of the process, with its terminal carboxylate group playing a critical role in effective inhibition. However, if the terminal carboxylate group is solely responsible for AOAA’s inhibitory effect, then other monoamine compounds containing a carboxylate group should exhibit similar inhibitory potency as AOAA.

### Significance of O1 atom in AOAA structure over its monoamine analogue 2- aminobutanedioic acid (Aspartic Acid/Asp)

To further verify if the terminal carboxylate of AOAA is solely responsible for inhibition efficacy. We selected a monoamine class analogue (non-reactive substrate) of AOAA, 2- aminobutanedioic acid, for further investigation where the terminal carboxylate was preserved (O1 atom deleted). Our in-silico investigation suggested that the Aspartic Acid (Asp) accommodated within the active core close to PLP **(Fig S6A)** and residues S72, N74, Y128, Y148, G181, G225, G227, E228 with a minimal backbone RMSD (3.5Å) and ligand RMSD (1.2-4.8Å) respectively as observed from (100ns) MD simulation **(Fig S6B)**. MD data suggested that the terminal carboxylate group of Asp could also provide interactions with T75 and Q147, respectively **(Fig S6C)**. Being a substrate class molecule, we further expected Asp to form external aldimine in *Mtb*CBS and also maintain pre-organized space in the active site as identified from PLP-Ser (PDB ID: 7XOY) and PLP-AOAA complexed (PDB ID: 9U7O) WT-*Mtb*CBS structures respectively. However, to identify external aldimine formation by Asp in *Mtb*CBS, we have selected the K44A mutant for this study to avoid confusion between internal and external aldimine signals. Though we have found an external aldimine while incubated with 2-aminobutanedioic acid/Asp (10mM) with 10μM K44A mutants, however, the extent of PLP-external aldimine formation was relatively less as compared with the canonical substrate L-Ser **(Fig S6D)**. We could not observe significant inhibition efficiency relative to 1mM AOAA-treated WT *Mtb*CBS, indicating that Asp could not form a stable complex with PLP. Both the substrate (L-Cys) and K44 likely destabilized the PLP-Asp complex. This led us to conclude that hydroxylamine-based compounds are more effective inhibitors than their monoamine counterparts (2-aminobutanedioic acid). Finally, we extended our hypothesis if the terminal carboxylate of AOAA interaction with polar residues T75 or Q147 is only responsible for the observed inhibition with AOAA or interaction with any other non-polar residues could also exert a similar stabilizing effect on Inhibitor candidates.

### Molecular insights for functional infidelity of O-Benzylhydroxylamine (PhCH2ONH2) in WT-MtbCBS

Targeting the reactive side chain and inactivating the cofactor is a common strategy for designing inhibitors against enzymes ^29,30^. If neutralizing the reactive side chains (T75 and Q147) or potentially locking PLP using hydroxylamine or a synergic approach influences the inhibitor efficiency, we selected O-Benzylhydroxylamine (used for inhibitor design against Aminotransferase PDB-3DTG and as an antimalarial compound ^31^ PDB-7ZEA) that contained a bulky phenyl group, which can potentially interact with other hydrophobic residues (instead of polar T75and Q147) in the active pocket. Our MD simulation data suggested that PhCH2ONH2 could interact with several hydrophobic residues, including P70, Y128, Y129, and Y148 (more strongly than MOA **(Fig 5A, S7A)** in the active pocket. Additionally, the extent of aldimine drop and H2S production relative to AOAA suggested the reversible nature and a highly compromised inhibitor efficiency of PhCH2ONH2 in the presence of substrates **(Fig 5B)**. Therefore, to decode the factors behind poor inhibition efficacy, we performed Cryo-EM to reveal the molecular insights of PhCH2ONH2 in WT-*Mtb*CBS. Similar to the AOAA-bound WT-*Mtb*CBS Cryo-EM structure, the tetrameric geometry of the enzyme was maintained with a minimum backbone structural deviation in each subunit **(Fig 5C)**. From our cryo-EM density map, we could not find any strong density close to PLP that could suggest the formation of a PhCH2ONH2-PLP complex. Our reconstituted atomic model (PDB ID: 9U7N) projected a strong EM density over the PLP-44Lys- ζ NH2 internal aldimine complex **(Fig 5C, Movie-2)**. The lack of strong density for the PhCH2ONH2-PLP complex could reflect a weak binding affinity of PhCH2ONH2 towards active site PLP where internal base 44Lys-ζ NH2 acts as a strong nucleophilic competitor. Interestingly, we found a strong extra density (fitted for PhCH2ONH2 ligand) close to Y128, Y129, and Y148 residues as initially predicted from our in-silico investigation. From our atomic model, the prior location of this molecule, from cofactor PLP-K44 complex and catalytically active T75 & Q147, was depicted **(Fig 5C)**. However, more precisely, we could find its conformation at the entrance of the active site channel where Y128, Y129, and Y148 could act as gatekeeper residues ^32^. However, the ability of the PhCH2ONH2 molecule to act like a classical channel blocker was not reflected in the inhibition studies with *Mtb*CBS. To explore why hydroxylamines function as superior PLP inhibitors, we conducted QM/MM investigations.

**Figure 5.**
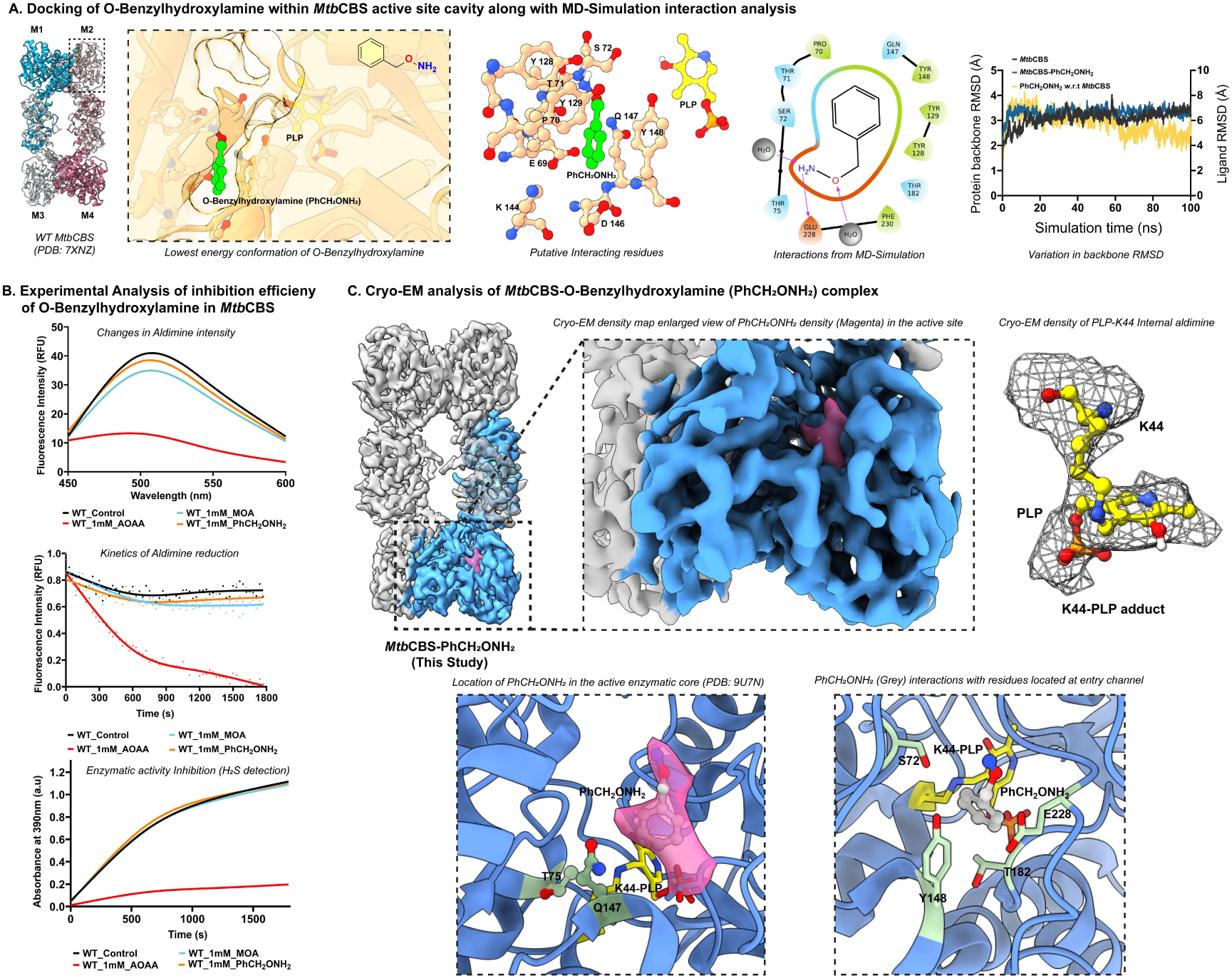
*Structural understanding of inhibition infidelity of O-Benzylhydroxylamine*. ***A.*** In-silico molecular docking analysis to predict lowest energy conformation of MtbCBS-O-Benzylhydroxylamine ligand and the putative interacting residues. The tetramer of MtbCBS with the lowest energy docked conformation of ligand coloured showing individual protomeric units. An enlarged view of the M2 protomer is marked by a black dashed rectangle. Ligand, close proximal residues, and cofactor PLP were rendered in a ball and stick representation on the transparent protein surface. Ligand was accommodated in a cavity away from PLP. Putative ligand interacting residues in 3D orientation. The 2D-ligand interaction diagram from the MD simulation was depicted. Ligand is coloured lime green, and PLP is coloured yellow. Protein backbone RMSD changes without a docked ligand (black) in the presence of the PhCH2ONH2 ligand (blue), and the ligand RMSD changes (yellow) over the time course of MD simulation. ***B.*** Line plots showing the fluorescence emission spectrum from 450-600nm recorded after 30 minutes of a kinetic study showing the extent of aldimine species present, kinetics of aldimine changes (λex/em=410/510nm), and the kinetics of H2S production inferred through lead acetate- assay in MtbCBS treated with 1mM of inhibitor AOAA and molecular mimics (Control/Untreated: black, AOAA: red, MOA: light blue, PhCH2ONH2: light brown). ***C.*** Cryo-EM analysis of MtbCBS-O-Benzylhydroxylamine complex. Cryo-EM density map (EMD-63940) showing the intact tetrameric arrangement of O-Benzylhydroxylamine bound MtbCBS (Single protomeric subunit coloured blue). Enlarged view showing the cryo-EM density of the bound O-Benzylhydroxylamine (dark pink). The corresponding cryo-EM density for PLP-K44 in the active site cavity is rendered as mesh. Enlarged views of the atomic model depicting the bound O-Benzylhydroxylamine (density coloured pink; PDB ID 9U7N) at the entrance of the active channel and the other view indicating the proximal interacting residues with O-Benzylhydroxylamine.

### QM/MM investigations

To investigate the factors contributing to the increased stability of PLP-hydroxylamine adduct over monoamines, we carried out QM/MM analysis to determine the nucleophilicity of the molecules (AOAA, MOA, and 2-aminobutanedionic acid), reactivity of the molecules and stability differences in PLP-molecule adduct. The molecular orbitals of the geometry-optimized compounds and PLP-molecule adducts were visualized through multiwfn **(Fig 6A- B)** ^33^. The energy difference between HOMO and LUMO orbitals is highest for MOA (7.898 eV) relative to other compounds, suggesting unfavourable electron transfer, making it highly reactive. This could be due to the absence of multiple reactive functional groups in MOA compared with other molecules. The energy gap decreases from MOA (7.898 eV) > AOAA (6.688 eV) > Asp (6.513 eV) > PhCH2ONH2 (6.412) **(Fig-6C)**. We find that Aspartic acid exhibited the least chemical potential (-4.274 eV), highest dipole moment (5.184 debye) and lowest electronegativity (-2.805 eV) relative to other hydroxylamine class compounds from the trends depicted **(Fig-6C)**. This could indicate that Aspartic acid, an amino acid-like substrate, cannot be a strong reacting nucleophile. All the other molecules showed better nucleophilicity and better chemical potential to react than aspartic acid. Molecular properties were computed in a similar fashion for PLP-molecule adducts **(Fig 6D)**. We observe that the PLP-AOAA adduct showed the least chemical potential (-4.697 eV), highest dipole moment (9.463 debye), and low energy gap (4.491 eV) making it the most stable PLP adduct among the studied PLP-molecule adducts. Other molecular properties, such as maximum electronic charge, polarizability, and nucleophilicity, were also computed **(Table-4)**. We infer that all hydroxylamine class molecules showed better reactivity than the selected monoamine aspartic acid. The presence of an extra oxygen atom could be the additional factor contributing to the increased reactivity of hydroxylamines class molecules. Therefore, we conclude that the synergically binding to PLP and locking specifically to reactive side chain residues boosted PLP-oxime stability and projected AOAA as a potent CBS inhibitor.

**Figure 6.**
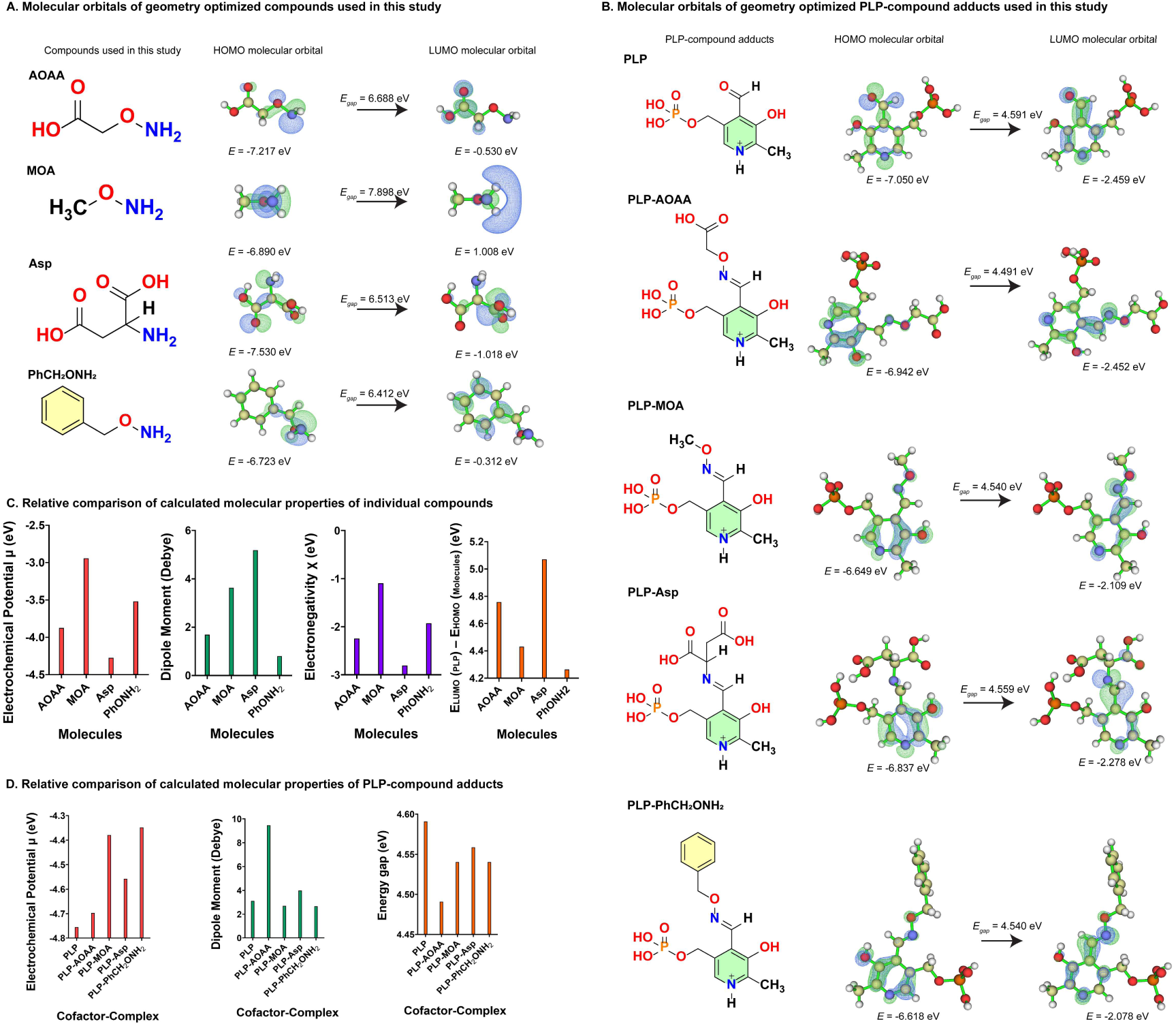
*Quantum Mechanical (QM) investigations*. ***A.*** Molecular orbitals of geometry optimized compounds (AOAA, MOA, Aspartic Acid, O-Benzylhydroxylamine). The optimized geometries of the compounds mentioned were determined at B3LYP/6-311G(d,p) level in the ground state. The chemical structures were drawn using ChemDraw. The HOMO and LUMO of the compounds visualized using multiwfn and their corresponding energies, HOMO-LUMO energy difference were indicated for each case (energy values in units of eV). ***B.*** The multiwfn visualization of the HOMO, LUMO for the geometry-optimized PLP- compound adducts (PLP-AOAA, PLP-MOA, PLP-Asp, PLP-PhCH2ONH2) along with the control PLP cofactor. The calculated energies of HOMO, LUMO and the energy gap between the orbitals. ***C.*** Bar plots show the relative differences in chemical potential, dipole moment, electronegativity, and the energy difference E(LUMO) of PLP and E(HOMO) of the compound. ***D.*** Bar plots indicating the relative differences in chemical potential, dipole moment and energy gap of PLP-compound adducts.

## Discussion

The in-silico approach has been widely accepted for structure-based drug design (SBDD) ^34^. However, in the field of medicinal chemistry of enzymes, synthetic electrophilic molecules were designed purposefully based on the understanding of reaction mechanisms ^35,36^. Besides targeting cofactors, locking active side chains could provide an alternative strategy for targeted drug design for metabolic enzymes ^37^. Therefore, synthetic electrophiles that specifically target the critical nucleophilic side chains taking part in the critical step of the reaction could accelerate the process of development of new therapeutics. Irreversible chemical modification to the reactive side chains, especially Cys, Lys, and Tyr, had been a key targeted strategy for inhibitor synthesis for several enzymes ^38–40^. Drugs like Osimertinib, Sotorabsib, Salicylaldehyde derivatives and Propargyl glycine were predicted to react with cysteines and catalytic lysine, forming stable imines, thereby inhibiting enzymatic activity in Hsp90, and CGL, respectively ^41,42^. Therefore, this approach is more efficient for designing better drug candidates, while the conventional structure-based drug design is still effective for membrane proteins, including GPCR, ion channels and other cellular target proteins ^43,44^. The limited structural insights of holo-CBS enzymes and their intermediates (aldimine, carbanion, amino acrylate intermediate) were restricted only to the cofactor PLP centre ^45–48^. Though AOAA interaction with *h*CBS and other PLP-dependent enzymes (β-phenylalanine aminotransferase, CSE, γ-aminobutyrate aminotransferase, human kynurenine aminotransferase) were investigated earlier, the mechanistic aspects of enzymatic inhibition were never investigated in depth except highlighting the AOAA-PLP covalent/non-covalent bonding ^49–53^. Our in-silico and QM/MM calculations reflected that the nucleophilic hydroxyl amines could effectively stabilize the free PLP complex compared to its monoamine candidates. Therefore, the old hypothesis of AOAA non-specifically targeting other PLP-based enzymes still remains true ^54–56^. However, its selectivity (irreversibility) towards PLP-dependent enzymes was controlled by other essential factors in the context of the enzyme and its active catalytic core. Being hydroxyl amine class molecules that were capable of forming the PLP-Oxime complex, the poor inhibitor efficacy of MOA and PhCH2ONH2 suggested that the PLP-adducts required additional stabilization inside the active core of the protein. Such additional stabilizing interactions were observed for PLP-AOAA adduct with active side chains thereby further boosting the inhibition potency of AOAA towards the WT-*Mtb*CBS. In all other cases, PLP- MOA, PLP-Asp and PLP-PhCH2ONH2 reside in the active site via weak interaction with other non-specific side chains or block the active channel, making these hydroxylamine compounds less effective than AOAA, respectively. Our understanding was further consolidated when we proved the functional neutrality (functionally inactivity) of AOAA towards the T75A and Q147 variants. Therefore, the synergic efforts of AOAA for *Mtb*CBS that reflected its functional potency over other hydroxyl amine and mono-amine analogues could also explain its (inhibition) efficacy for other structurally conserved CBS enzymes and will open new insights for elucidating and discovering PLP-targeting and non-PLP-targeting inhibitors in the field of medicinal chemistry.

## METHODS

### 1. Cloning, Expression and Purification of recombinant WT MtbCBS and mutant variants

*Mtb*CBS Wild type and mutant variants were expressed in *E. coli* BL21 (DE3) cells and purified as mentioned previously ^22^. Plasmid containing the wild-type *Mtb*CBS gene was used as the template for introducing site-directed mutations (SDM). The following mutations, T75A, Q147A, Q147Y, Q147N and Q147E, were introduced in non-overlapping primers. Primer details used for generating mutants are provided in ***Table-3***. Briefly, the template was PCR amplified with the mutation primers using Q5 DNA polymerase (10 units/ml) (NEB). SDM was performed by digesting the template by incubation with DpnI (800 units/ml) (NEB) for 4 hours, and the digested template was removed through agarose gel electrophoresis. The mutant DNA was gel-extracted, and the DNA concentration was measured using NanoDrop 2000c (Thermofisher Scientific). DNA sample (10 ng) was incubated with T4-PNK (500 units/ml) (NEB) and T4-DNA ligase (20,000 units/ml) (NEB) overnight at 16°C and finally transformed into *E. coli* TOP-10 competent cells through heat shock (***Table-2***). The transformed bacterial cells were screened using the same mutant primers through colony PCR, and the positive colonies were used to prepare primary culture (10 ml of LB supplemented with Kanamycin) for plasmid isolation. The plasmids were isolated from the bacterial cells, and the mutation was further confirmed by DNA Sanger sequencing. We proceeded with two-step purification Immobilized-Metal Ion Affinity Chromatography (IMAC/Ni-NTA) (Qiagen) followed by Size Exclusion Chromatography (AKTA-Pure, GE Healthcare) in gel-filtration buffer (50mM Tris, 150mM NaCl, pH 7.5) for both WT *Mtb*CBS and its mutant variants. In all purifications, recombinant proteins were eluted at 14ml volume corresponding to a tetrameric molecular weight ∼200kDa (Superose-6 Increase 10/30 GL, Cytiva). The purity of recombinant enzymes was assessed through SDS-PAGE **(Figure S2A)**. All the reagents used for this work are mentioned in ***Table-5***.

### 2. Enzymatic assay

Enzymatic activity of WT *Mtb*CBS and its mutant variants were monitored using lead-acetate- based H2S (By-product from β-elimination reaction) detection ^10^. All the in-vitro enzymatic assays were performed using freshly purified CBS enzymes (WT/mutants) with a final protein concentration of 10μM. Briefly, recombinant *Mtb*CBS (WT and mutants) was incubated with varying working concentrations of AOAA (100μM**-**1mM) in reaction buffer (50mM Tris, 150mM NaCl, pH 7.5) for 1 minute before adding 50 mM L-cysteine to the reaction mixture **(Figure S3C)**. Lead acetate solution (4mM) was added so that the total reaction volume became 200μL in different wells of a 96-well tissue culture plate (Tarsons) under closed conditions for non-kinetic experiments at 25°C. Steady-state kinetics was done by mixing WT *Mtb*CBS with SAM (0.5mM, in SAM-treated sets) followed by L-cysteine (0-80mM) and incubated for 1 min at 25°C **(S2C)** prior to lead acetate (4mM) addition. For assessing mutant enzymatic activity, *Mtb*CBS (WT/mutants) were incubated with three different AOAA concentrations (10μM, 100μM, 1000μM) for 2-3 mins at 25°C, then L-cysteine (50mM) and lead acetate (4mM) were mixed to a final reaction volume of 200μL **(Figure 3B&4B)**. Activity inhibition by 2- aminobutanedioic acid is conducted by incubating WT *Mtb*CBS with 2-aminobutanedioic acid (10mM) for 1 min, followed by L-cysteine (50mM) **(Figure S6D)**. The extent of H2S production was measured using the plate-based lead acetate assay (T=25°C, λ=390nm) on a Varioskan plate reader with a 30-second time interval for a total measurement time of 30 minutes in kinetic experiments. Enzymatic assays (Kinetics) were performed in three independent sets. The mean values were plotted as bar plots and curve plots using GraphPad Prism (https://www.graphpad.com/).

### 3. Quantification of PLP-External Aldimine and PLP-Oxime species

The identification of PLP-external aldimine and PLP-oxime species and their quantification were made using a fluorescence-based detection assay. The assays were carried out in 50 mM Tris (pH 7.5) and 150 mM NaCl reaction buffer. The AOAA, L-serine, and L-cysteine stock solutions were prepared with the same reaction buffer. All reactions were conducted at 25°C with a total reaction volume of 200μL and a final enzyme concentration (WT/mutants) of 10μM. For the CBS-AOAA interaction study, the reaction was prepared by incubating *Mtb*CBS (WT/mutants) with increasing concentration of AOAA (20-200μM) for 10 mins **(Figure S3A- B)**. AOAA-PLP-external aldimine competition assay was done by initially incubating the WT *Mtb*CBS with 10mM Substrate (L-serine/L-cysteine) for 10 mins, then different concentrations of AOAA (5-1000μM) were added and incubated for 2 mins prior to detection **(Figure S3D- E)**. Suicidal inhibition assay was carried out by incubating WT *Mtb*CBS with increasing AOAA concentrations (20-200μM) for 2 mins, then 10 mM of the substrate (L-serine/L-cysteine) was added to the reaction solution and further incubated for 10 mins **(Figure 2A)**. IC50 concentrations of AOAA were determined by treating WT *Mtb*CBS with 10mM substrates (L-serine/L-cysteine) for 10mins followed by the addition of AOAA (5-5000μM) and incubating for 1-2 mins **(Figure S3F)**. AOAA binding to *Mtb*CBS mutants was studied by pre-incubating *Mtb*CBS (WT/T75A/Q147A) with L-serine (10mM) for 10 mins followed by three different AOAA concentrations (10μM, 100μM, 1000μM) for 2 mins **(Figure 3C&3E)**. The kinetic assay was started by mixing AOAA (1mM) with *Mtb*CBS (WT/Mutants) pretreated with 10mM L-serine for 1 min **(Figure 3F)** in a 96-well assay plate and analysed on a plate reader. Aldimine reduction and oxime formation were measured kinetically by exciting at wavelengths 410nm and 350nm and recording the fluorescence emission intensities at 510nm and 450nm, respectively. Fluorescence measurements were carried out in kinetic mode for 30 minutes in a Varioskan Flash plate reader with a bottom reading (Bandwidth 5nm, PMT voltage set to medium). The mean values were plotted using GraphPad Prism.

### 4. Negative staining TEM analysis

Freshly purified *Mtb*CBS WT and mutants (SEC purified) were visualized under TEM to understand the oligomeric state and homogeneity of the protein. Untreated samples (WT, T75A and Q147A mutants), 1mM AOAA-treated WT and 1mM PhCH2ONH2-treated WT *Mtb*CBS (10μM) were diluted to a final protein concentration of 0.05 mg/ml with PBS (1X) for TEM analysis. 3μL of the diluted sample was applied onto a freshly glow-discharged carbon-coated Cu grid (PELCO) for 30 seconds (20mA maintained for 30 seconds with a glow time of 10 seconds under atmospheric air conditions). Further, the excess sample was removed by blotting, and staining was performed using a 1% uranyl acetate solution. Samples were visualized under (Talos L120C) electron microscope equipped with a CETA camera with a nominal magnification of 92kX (Calibrated pixel size of 1.52Å). In order to understand the geometrical states of CBS enzymes, we performed reference-free 2D classification for Untreated WT, AOAA-treated WT, and PhCH2ONH2-treated WT separately using the EMAN-2.9 software package ^57^. In all cases, we find that the tetrameric arrangement of CBS remains intact with and without inhibitor treatment **(Figure S2B, S2D & S8C)**.

### 5. Cryo-EM sample preparation

Graphene oxide-coated Quantifoil grids (EMS, Cu 300 mesh, R1.2/1.3) were used for cryo-EM sample preparation for *Mtb*CBS-AOAA samples. Graphene oxide grids were prepared following the protocol mentioned ^58^. AOAA-*Mtb*CBS complex was prepared by gently mixing freshly purified WT *Mtb*CBS (1mg/ml) with AOAA (1mM) for 1 min on ice (4-10°C) with a protein to AOAA ratio of 1:50. 4μL of undiluted AOAA incubated *Mtb*CBS sample was applied to the prepared graphene oxide-coated quantifoil grids, and blotting was carried out using Vitrobot Mark-IV (blot time 8.5s, without additional blot force). The samples were plunge-frozen in liquid ethane (-183°C) and stored under liquid nitrogen conditions (-196°C).

PhCH2ONH2-*Mtb*CBS was prepared in the same way as the AOAA sample, resulting in the final concentration of 1mg/ml and 1 mM for *Mtb*CBS and PhCH2ONH2. The sample was incubated on ice for 10 minutes with continuous pipetting. 4μL of this freshly prepared sample was applied to the carbon-coated quantifoil grids (EMS/TedPella, Cu 300 Mesh, R1.2/1.3) under controlled conditions inside the Vitrobot Mark-IV system. Blotting was carried out with the same parameters mentioned above, and the plunge was frozen in liquid ethane.

### 6. Cryo-EM data acquisition

The cryo-EM data was collected using a Talos Arctica 200keV electron microscope (Thermofisher Scientific) equipped with a K2 Summit Direct electron detector (GATAN Inc). 5,980 movies were recorded for the *Mtb*CBS-AOAA sample through automatic data collection using Latitude-S software (GATAN) in which the total electron dose of 45e^°^/Å^2^ at the specimen level was fractionated among the 20 frames with defocus varying between -1.25μm to -2.25μm at a nominal magnification of 54,000X corresponding to a calibrated pixel size of 0.92Å respectively ^59^. For the *Mtb*CBS-PhCH2ONH2 sample, 5,346 movies were recorded with similar data collection parameters (***Table-1***).

### 7. Cryo-EM data processing

The cryo-EM data processing pipeline followed for the AOAA-treated *Mtb*CBS dataset was provided in **Figure S4A**. The collected 5,980 movie files of AOAA incubated *Mtb*CBS were processed using cryosparc version v4.5.3 ^60^. Patch motion correction and CTF estimation were carried out, and micrographs with CTF fit resolution of less than 7Å were selected. 5,675 micrographs curated based on the above fit-resolution. Particles were picked using blob-picker. Around ∼10 lakh particles picked were carefully inspected. The particles were extracted using a box size of 292 pixels, and multiple rounds of 2D classification were performed. 2D class averages with better features and better Fourier Ring Correlation (FRC) were taken for heterogeneous classification. The heterogeneous classification was done to classify ∼6.6 lakh particles into nine 3D classes (D2 symmetry) using a previously imported 40Å low-pass filtered map (PDB 7XNZ) as the initial reference ^22^. 3D-class with better resolution and a decent particle number was further refined non-uniformly (NU-Refinement) using a soft solvent mask (generated based on the threshold from UCSF ChimeraX ^61^/Dynamic Mask) to a 4.1 Å resolution map. The map resolution was further improved by reference motion correction, global CTF refinement and local refinement to achieve a final cryo-EM map resolution of 3.96 Å. The validation plots (GSFSC, Angular distribution, map-model validation), local resolution estimate, and the fitting of the cryo-EM density map to the atomic model, along with the PLP- AOAA adduct density, were provided in **Figure S4B-G.** The cryo-EM data processing pipeline **(Figure S5A)** of *Mtb*CBS treated with PhCH2ONH2 dataset was pre-processed (Motion corrected, Patch CTF estimated with 5,346 micrographs), yielding ∼5,230 micrographs with ∼8 lakh particles. The particles were classified using 2D classification, and particles belonging to better 2D class averages (FRC & particle number) were classified heterogeneously into five 3D classes (C1 symmetry) using the above-mentioned 40 Å low-pass filtered map as a reference. NU refinement using particles from classes (4&5) further improved class 4, showing better features and resolution. Particle quality was enhanced by reference-based motion correction and global CTF refinement jobs. Local refinement using the polished particles brought down a resolution to 4.19 Å. The validation plots and local resolution estimate were provided in **Figure S5B-E**. The density for the bound PhCH2ONH2, along with the fitting with the atomic model, was given in **Figure S5F-G.** The data collection and processing statistics are provided in ***Table-1***.

### 8. Cryo-EM model building, refinement, and validation

The available PDB for *Mtb*CBS (PDB ID: 7XNZ) was used as the starting model, and rigid-body fitting into the obtained cryo-EM maps was performed in UCSF ChimeraX ^61^. Real-space refinement of the starting model 7XNZ was performed using PHENIX (phenix:real_space_refine) ^62^. The refined model and the corresponding cryo-EM maps were imported in COOT to manually fit and inspect the refined model ^63^. The coordinates of the PLP-AOAA (IK2) adduct were obtained from PDB and were fitted manually in the unmodelled density at the active site pocket using COOT ^63^. The model with fitted PLP-AOAA adduct was iteratively improved through phenix:real_space_refine and manual modifications using COOT^63^. A similar approach was followed for generating the atomic model for the PhCH2ONH2 bound *Mtb*CBS structure. The PDB structure of O-Benzylhydroxylamine (OBZ) was docked in the unmodelled density in the cryo-EM map of PhCH2ONH2-treated *Mtb*CBS maps, and the resulting atomic model was refined using PHENIX ^62^. The validation was carried out using phenix:validation_tool in PHENIX, and the obtained model validation scores were provided in ***Table-1***.

### 9. Molecular docking analysis

Docking of 2-aminobutanedioic acid (C4H7NO4), methoxyamine (CH3ONH2), and O-benzylhydroxylamine (PhCH2ONH2) against the available PDB structure of WT *Mtb*CBS (PDB ID 7XNZ) was carried out separately using the SwissDock server ^64^. The ligand structural restraint files were fetched from the PubChem library in 3D SDF format. The ligands were optimized (geometry and energy) based on the universal force field UFF through the Steepest Descent algorithm using the Avogadro (an open-source molecular builder and visualization tool) ^65^. Rigid-body docking was carried out by the Vina method in which a cubic box of dimension 30Åx30Åx30Å was centred around the active site region of one of the subunits of *Mtb*CBS (140.0, 85.0, 144.0) and docking was carried out with a sampling exhaustivity of 8. Subsequently, 20 different docking conformations were visualised using UCSF chimeraX, and the conformation with the lowest binding energy was selected for MD simulation analysis ^61^. The interacting residues in the lowest binding energy conformation were visualised using Discovery Studio, and all the representative figures were made using UCSF ChimeraX ^61^.

### 10. MD simulation analysis

The MD simulation studies used the Schrödinger Maestro molecular modelling package (Schrödinger Release 2024-4: Maestro, Schrödinger, LLC, New York, NY, 2024). Since the docking was performed using rigid receptor mode and MD simulation was carried out to assess the ligand-induced conformational changes in the active site of *Mtb*CBS, the resident time of the ligand in the active site. The obtained best binding pose was imported into the Maestro suite, and the protein preparation workflow was followed to optimize the protein structure and detect the docked ligand. A standard protonation state at pH 7.5 was used, and the hydrogen atoms were incorporated. The system for simulation was prepared using the Desmond system builder, in which the entire protein was placed in an orthorhombic box and solvated using single point charge (SPC) solvent molecules. The charge on the protein was neutralised by adding sodium ions. Around 0.15 M of NaCl was added to the system to maintain the physiological conditions. The systems were described using the OPLS_2005 forcefield (version 64135). The simulation system was relaxed following Desmond’s default eight-stage relaxation protocol. The isotropic Martyna-Tobias-Klein barostat maintained pressure at 1 atm, and the Nose-Hoover thermostat was used to maintain the temperature at 300 K. The Smooth particle mesh Ewald method was used to evaluate long-range coulombic interactions with a short-range cutoff of 9.0 Å. The simulation (Production MD) was carried out using the Desmond MD simulation module using an NPT ensemble for 100 ns. The simulation was analysed using the simulation interaction diagram module.

### 11. Density Functional Theory (DFT) calculations

All computational calculations were performed with the Gaussian 16 using density functional theory (DFT) ^66^. The Avogadro was used to obtain the molecule’s initial cartesian coordinates^65^. The molecular geometries of all molecules utilised in this work were optimised through density functional theory calculations, using the FREQ and RB3LYP method calculations with 6-311G (d,p) as the basis set. The resulting optimized structures were used to compute the molecular orbitals using DFT calculations. The obtained orbitals of the molecules were rendered using multiwfn ^33^. Other parameters like Energy gap, chemical potential, and nucleophilicity were computed (***Table-4***).

## Acknowledgements

The authors acknowledge the Advanced Center for Cryo-Electron Microscopy facility, Negative Staining TEM facility and Molecular Biophysics Unit Central Instrumentation facility for performing the experiments. We also thank the Division of Biological Science, HPC-Beagle, for processing Cryo-EM data and SERC-RNC for carrying out quantum mechanical calculations. We thank Mr Drose Inku Shane, Ms Keerthana, Mrs Aparna Asok and Mr Madan Reddy for helping us collect data from the Central Instrumentation facilities. We acknowledge the Science and Engineering Research Board (CRG/2022/002674 & STR/2022/000006) for financial support. We also acknowledge the Department of Biotechnology DBT-BUILDER Program (BT/INF/22/SP22844 /2017), Department of Science and Technology DST-FIST (SR/FST/LSII-039/2015), India, for funding the Advanced Center for Cryo-Electron Microscopy (ACCEM) facility at IISc-Bangalore. SP acknowledges the Prime Minister Research Fellowship for his fellowship (TF/PMRF-21-1313.03). AR acknowledges the financial support from the Ministry of Education (MOE) (STARS-1/171). BM acknowledges the Council of Scientific and Industrial Research (CSIR) for his fellowship. We acknowledge Mr Partho Pratim Das (PhD student, Molecular Biophysics Unit, IISc Bangalore) for the critical review of the manuscript.

## Author information

**Sainath Polepalli –** Molecular Biophysics Unit, Indian Institute of Science, CV Raman Avenue, Bangalore 560012, India. Email. **Anupam Roy -** Molecular Biophysics Unit, Indian Institute of Science, CV Raman Avenue, Bangalore 560012, India. Email.

**Bapan Mondal –** Molecular Biophysics Unit, Indian Institute of Science, CV Raman Avenue, Bangalore 560012, India. Email.

## Corresponding Authors

**\*Prof. Somnath Dutta -** Molecular Biophysics Unit, Indian Institute of Science, CV Raman Avenue, Bangalore 560012, India. Email.

**\*Prof. Amit Singh -** Centre for Infectious Disease Research (CIDR) & Department of Microbiology and Cell Biology (MCB), Indian Institute of Science, CV Raman Avenue, Bangalore 560012, India. Email.

## Authors contributions

SP and AR contributed equally to this work. SD, AS and AR Conceptualization, SP and AR performed protein purification, biophysical characterisation and biochemical studies. SP generated the mutant clones. SP, BM, and AR performed cryo-EM data processing. SP and BM carried out simulation and QM calculations. SD, AR, AS, SP, and BM analyzed the data. AR, SP wrote the manuscript and SD, AS, AR, SP and BM reviewed the manuscript. SD provided financial support for the work.

## Declaration

The authors declare no conflict of interest with the contents of this article.

## Data Availability Section

The final atomic model AOAA bound tetrameric *Mtb*CBS and refined cryo-EM map is available in wwPDB (PDB ID: 9U7O) and EMDB data with accession number EMD-63941. The atomic model and cryo-EM density map of PhCH2ONH2-bound *Mtb*CBS were also made available in the public domain (PDB ID: 9U7N, EMDB-63940). PDB accession IDs for native tetrameric form holo- *Mtb*CBS (PDB: 7XNZ) and *Mtb*CBS in the presence of S-adenosylmethionine and serine (PDB:7XOY) referred in this study had also been previously uploaded to wwPDB. All the supporting Figures (S1-S8) and Tables (1-5) are provided in the ***Supporting Information*** file.

---

**Table 1:** Cryo-EM data acquisition, data processing, model validation parameters.

| Parameters | CBS-AOAA | CBS-PhCH<sub>2</sub>ONH<sub>2</sub> |
| --- | --- | --- |
| Microscope type | Talos Arctica | Talos Arctica |
| Camera | K2 Summit DED | K2 Summit DED |
| Electron gun | Field emission gun | Field emission gun |
| Voltage | 200 kV | 200 kV |
| Electron dose | 45 e<sup>-</sup>/Å<sup>2</sup> | 46 e<sup>-</sup>/Å<sup>2</sup> |
| Dose per frame | 2.25 | 2.3 |
| Total number of frames | 20 | 20 |
| Apix value | 0.92 | 0.92 |
| Defocus range | -1.25 μm to -2.25 μm | -1.25 μm to -2.25 μm |
| Instrument used for plunge freezing | Vitrobot Mark IV | Vitrobot Mark IV |
| Data collection mode | Counting Mode | Counting Mode |
| Symmetry imposed | D2 | C1 |
| Number of movie files | 5,980 | 5,346 |
| Number of particles in the model | 171,221 | 216,091 |
| Map resolution | 3.96 | 4.19 |
| FSC threshold | 0.143 | 0.143 |
| Map sharpening B-factor (Å<sup>2</sup>) | -187.3 | -180.3 |
| MolProbity score | 3.17 | 3.04 |
| Poor rotamer (%) | 6.84 | 3.92 |
| Ramachandran plot |  |  |
| Favored (%) | 91.39 | 90.90 |
| Allowed (%) | 8.50 | 8.99 |
| Outliers (%) | 0.11 | 0.11 |

**Table 2:** Bacterial strains used in this study.

| Bacterial Strains used | Purpose |
| --- | --- |
| *Escherichia coli* BL21 (DE3) | Protein expression |
| *Escherichia coli* TOP-10 | Mutagenesis |

**Table 3:**
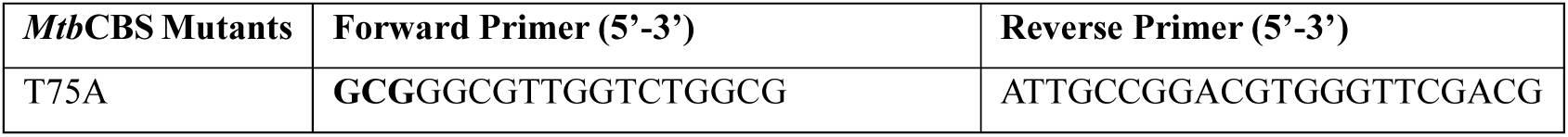
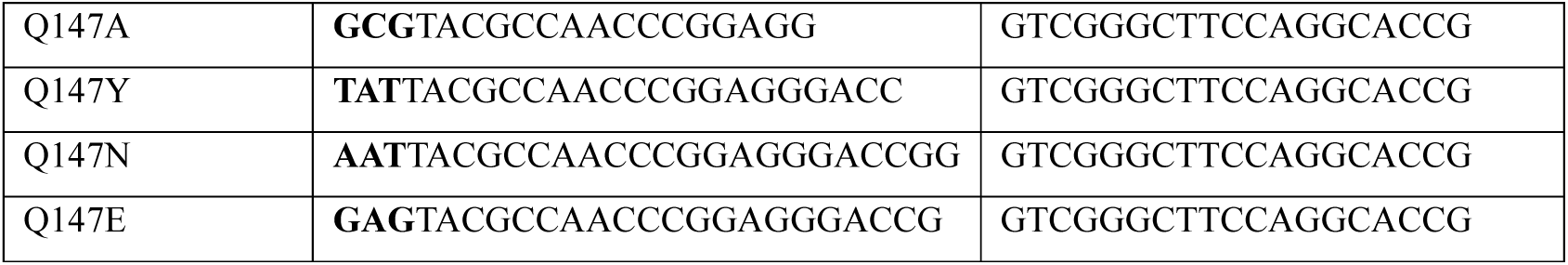
Primers used for performing Site-directed mutations.

**Table 4:**
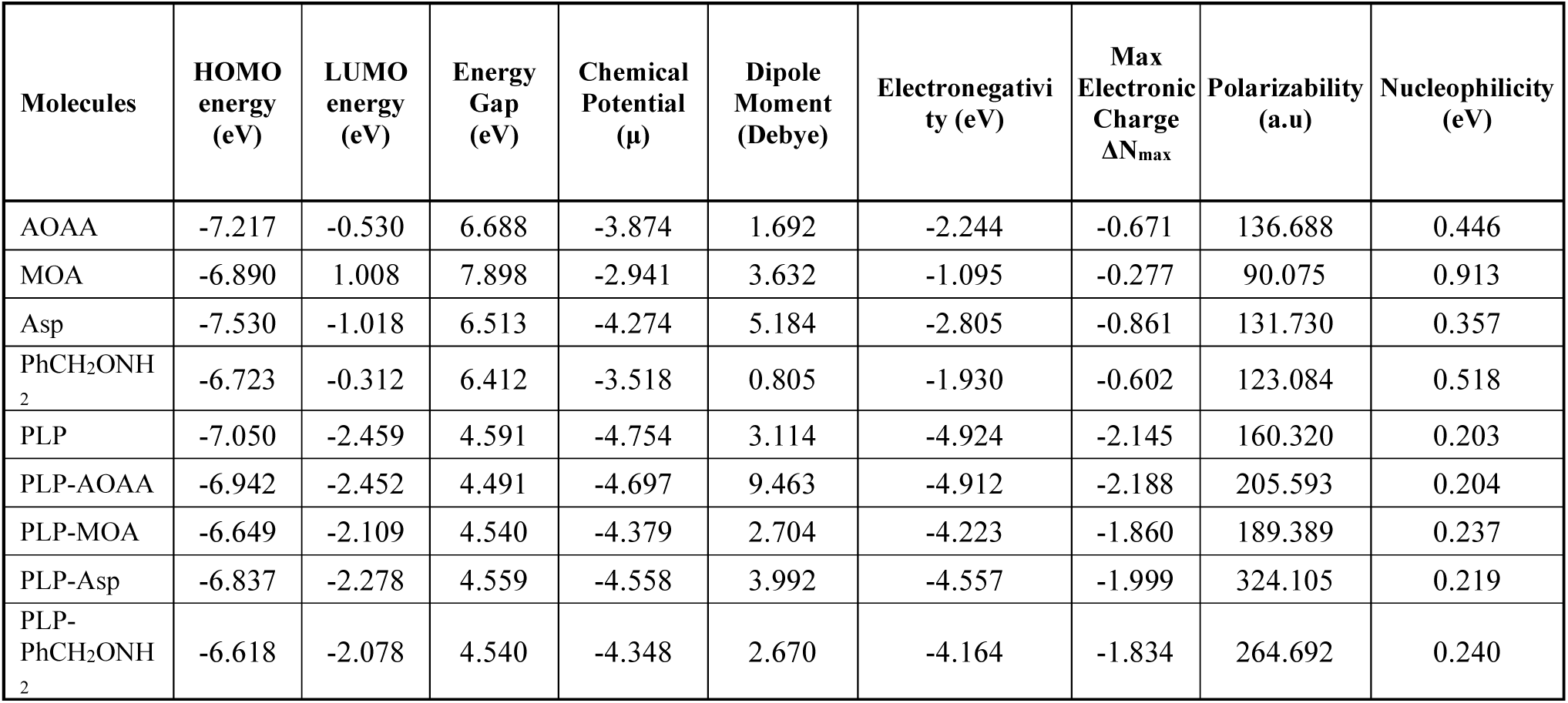
Computed properties of compounds and PLP-compound adducts.

| Molecules | HOMO energy (eV) | LUMO energy (eV) | Energy Gap (eV) | Chemical Potential ($\mu$) | Dipole Moment (Debye) | Electronegativity (eV) | Max Electronic Charge $\Delta N_{\max}$ | Polarizability (a.u) | Nucleophilicity (eV) |
| --- | --- | --- | --- | --- | --- | --- | --- | --- | --- |
| AOAA | -7.217 | -0.530 | 6.688 | -3.874 | 1.692 | -2.244 | -0.671 | 136.688 | 0.446 |
| MOA | -6.890 | 1.008 | 7.898 | -2.941 | 3.632 | -1.095 | -0.277 | 90.075 | 0.913 |
| Asp | -7.530 | -1.018 | 6.513 | -4.274 | 5.184 | -2.805 | -0.861 | 131.730 | 0.357 |
| PhCH<sub>2</sub>ONH<sub>2</sub> | -6.723 | -0.312 | 6.412 | -3.518 | 0.805 | -1.930 | -0.602 | 123.084 | 0.518 |
| PLP | -7.050 | -2.459 | 4.591 | -4.754 | 3.114 | -4.924 | -2.145 | 160.320 | 0.203 |
| PLP-AOAA | -6.942 | -2.452 | 4.491 | -4.697 | 9.463 | -4.912 | -2.188 | 205.593 | 0.204 |
| PLP-MOA | -6.649 | -2.109 | 4.540 | -4.379 | 2.704 | -4.223 | -1.860 | 189.389 | 0.237 |
| PLP-Asp | -6.837 | -2.278 | 4.559 | -4.558 | 3.992 | -4.557 | -1.999 | 324.105 | 0.219 |
| PLP-PhCH<sub>2</sub>ONH<sub>2</sub> | -6.618 | -2.078 | 4.540 | -4.348 | 2.670 | -4.164 | -1.834 | 264.692 | 0.240 |

**Table 5:** Chemicals and Reagents used in this study.

| Reagent | Source | Identifier |
| --- | --- | --- |
| Tris Base | Himedia | MB029 |
| Sodium Chloride | Himedia | MB023 |
| Potassium Chloride | Himedia | GRM3934 |
| Potassium di-hydrogen orthophosphate | Himedia | GRM3943 |
| Di-sodium hydrogen orthophosphate | Himedia | GRM1417 |
| Pyridoxal-5'-phosphate (PLP) | Himedia | RM1554 |
| Lysozyme | Sigma-Aldrich | 62971 |
| Kanamycin Sulfate | Himedia | MB105 |
| Phenyl Methyl Sulfonyl Fluoride (PMSF) | Himedia | TC706 |
| β-Mercaptoethanol | Himedia | MB041 |
| L-Serine | Sigma-Aldrich | 1077690010 |
| L-Cysteine | Sigma-Aldrich | 168149 |
| S-Adenosylmethionine hydrochloride (SAM) | Sigma-Aldrich | A7007 |
| Nuclease-free water | NEB | B1500S |
| Aminooxyacetic acid | MedChemExpress | HY-107994/CS-8158 |
| Methoxyamine | Sigma-Aldrich | 226904 |
| L-Aspartic Acid | Sigma-Aldrich | A9256 |
| O-Benzylhydroxylamine hydrochloride | Sigma-Aldrich | B22984 |
| Lead (II) Acetate trihydrate | Sigma-Aldrich | 32307 |
| Luria Broth | Himedia | M575 |
| Luria Bertani Agar Miller | Himedia | M1151 |
| Isopropyl β-D-1-thiogalactopyranoside (IPTG) | Himedia | RM2578 |
| Imidazole | Himedia | GRM1864 |
| Ni-NTA Agarose | Qiagen | 30230 |
| Coomassie Brilliant Blue R250 | Himedia | MB153 |
| Precision Plus Protein™ Dual Color Standards | BIO-RAD | 1610374 |
| Amicon Ultra-4 Centrifugal filter unit (30K MWCO) | Amicon | UFC9030 |
| Uranyl Acetate 98%, ACS Reagent | Polysciences, Inc. | 21447 |
| Superose 6 Increase 10/300 GL | Cytiva | 29091596 |
| Superdex 200 Increase 10/300 GL | Cytiva | 28990944 |
| EM grid (Carbon film 300 mesh, Copper) | Electron Microscopy Sciences | CF300-CU |
| EM grid (Quantifoil R12/1.3 300 mesh, Copper) | Electron Microscopy Sciences | Q3100CR1.3 |
| Sodium Dodecyl Sulphate | Affymetrix | 18220 |
| QIAprep Spin Miniprep | Qiagen | 27106 |
| QIAquick PCR Purification Kit for PCR Cleanup | Qiagen | 28104 |
| QIAquick Gel Extraction Kit | Qiagen | 28704 |
| Q5 High-Fidelity DNA Polymerase | NEB | M0491S |
| DpnI | NEB | R0176S |
| T4 Polynucleotide Kinase | NEB | M0201S |
| T4 DNA Ligase | NEB | M0202S |
| Ethidium Bromide | Himedia | MB071 |
| Ammonium persulphate | Himedia | MB003 |
| TEMED | Himedia | MB026 |
| Acrylamide | Himedia | GRM305 |
| N,N'-Methylenebis(acrylamide) | Himedia | MB005 |
| Glycine Hi-AR | Himedia | GRM199 |

