## Supporting Information for "Molecular insights into inhibitor action on a key bacterial metabolic enzyme *Cystathionine β-Synthase*"

### *Supporting Figures*

**A. Steps involved in catalytic cycle of *Mtb*CBS**

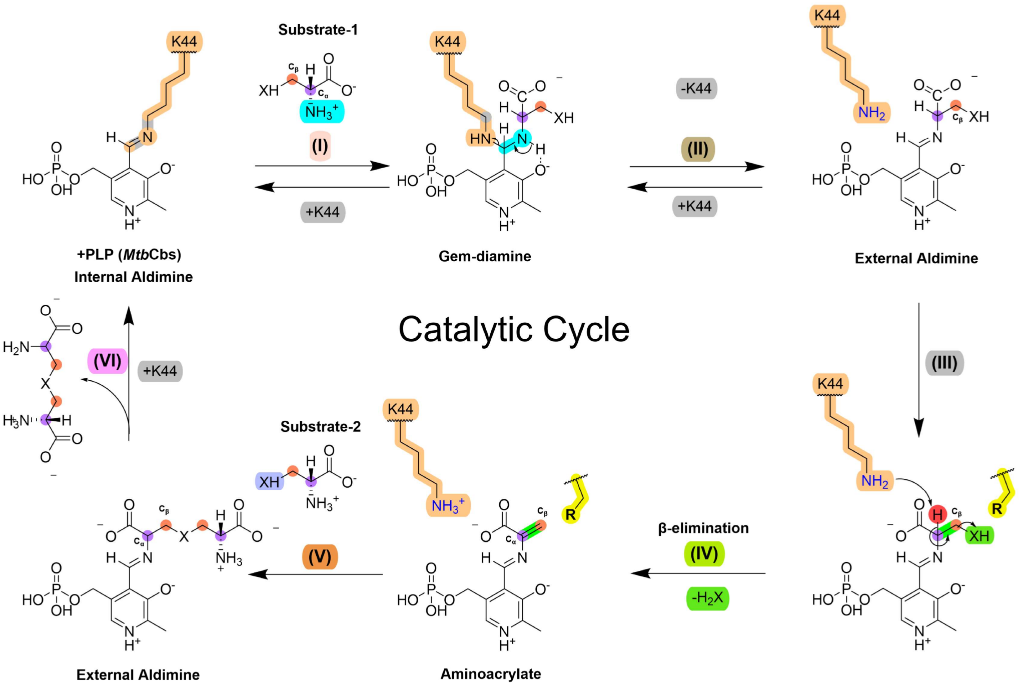


where X = O (L-Serine) or S (L-Cysteine)

**B. Active site core of CBS enzymes with conserved amino acid residues close proximity to PLP**

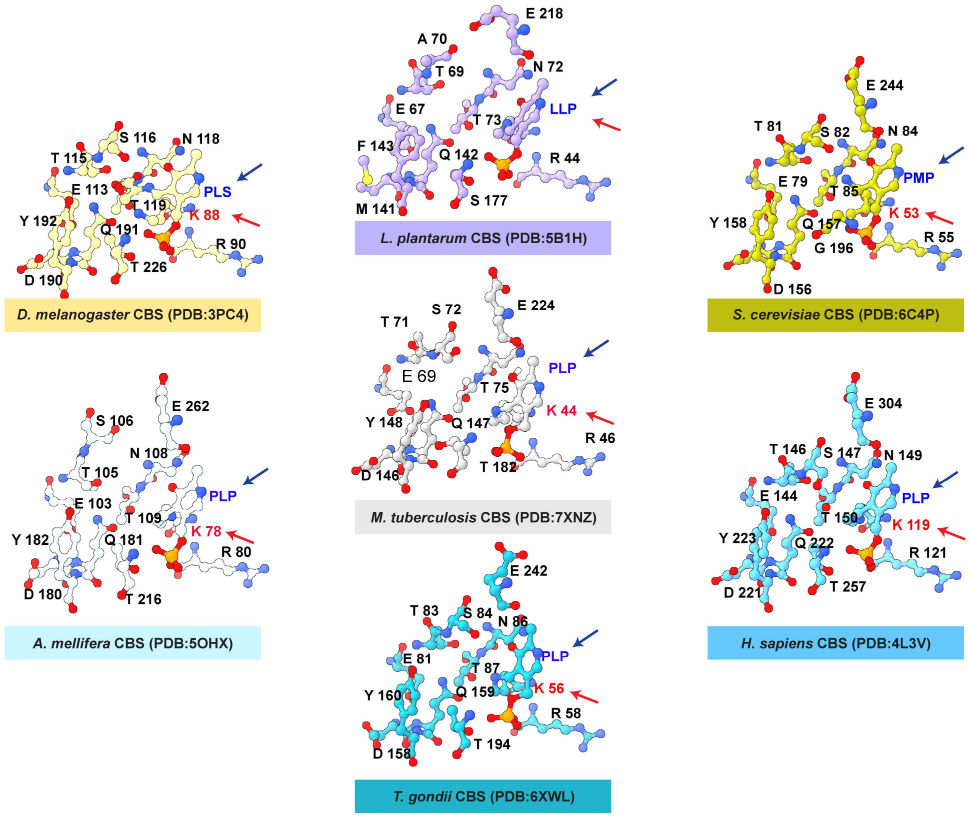

**Fig. S1. Steps involved in the catalytic cycle of *Mtb*CBS and conserved polar amino acid residues in representative CBS structures. A.** The entire catalytic cycle was divided into 6 major steps for easier understanding. Active site lysine residue is highlighted in light orange, the reactive amine group of substrate-1 is shown in Cyan and $C_\alpha$ & $C_\beta$ atoms of the substrate are marked with Purple and Salmon. The abstracted proton is coloured red, and the leaving group/ by-product released is coloured green. The reactive $C_\beta$ group in the second substrate molecule is highlighted in Melrose. The directionality of the reaction, along with the reversibility in K44 lysine's presence, is mentioned in all necessary steps. **B.** Active site core of *Mtb*CBS (*M. tuberculosis: **7XNZ***) showing the conserved amino acid residues in comparison with other crystal structures of CBS enzymes (*L. plantarum: **5B1H**; S. cerevisiae: **6C4P**; H. sapiens: **4L3V**; T. gondii: **6XWL**; A. mellifera: **5OHX**; D. melanogaster: **3PC4***) from other species respectively. The conserved residues were indicated in black. The PLP and catalytic lysine were marked with blue and red arrowheads.

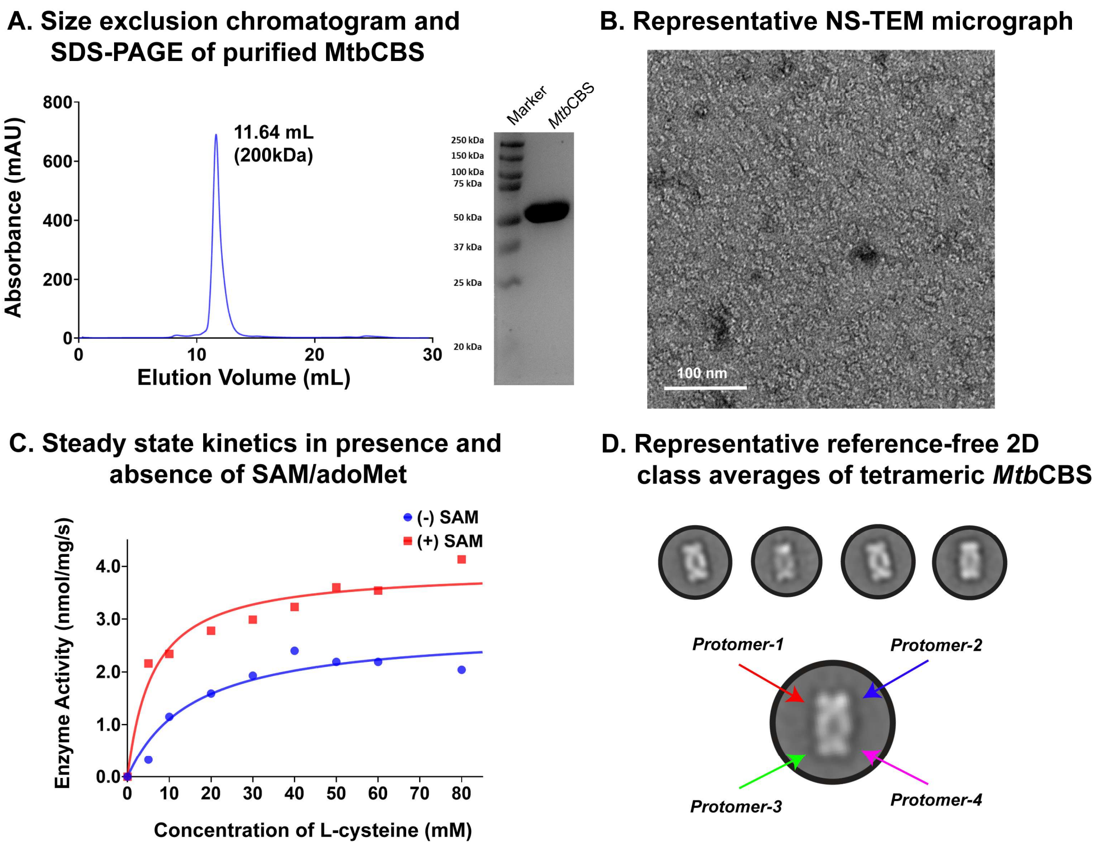


**Fig. S2. Purification of functionally active recombinant tetrameric *Mtb*CBS. A.** SEC profile of Ni-NTA purified *Mtb*CBS (mAU, milli-arbitrary units) obtained using Superose-6 HiLoad column; SDS-PAGE gel showing the SEC-purified *Mtb*CBS (Lane 2) along with the protein molecular weight marker (Lane 1). **B.** Representative Negative Staining (NS-TEM) micrograph of the SEC purified *Mtb*CBS showing isolated tetrameric orientations. **C.** Steady-state kinetics of SEC-purified *Mtb*CBS showing enzymatic activity dependence on L-Cysteine concentration (0-80mM) and the activity was found to enhance by ~2 fold in the presence of 0.5mM SAM (Red) relative to the control sample (No SAM). The calculated Kinetic parameters for *Mtb*CBS in the presence (and absence) of SAM were found to be $K_m$ 6.046mM (15.47mM) and $V_{max}$ 3.937nmol/mg/s (2.807nmol/mg/s). **D.** Representative reference-free 2D class averages of *Mtb*CBS, obtained using EMAN, showing the tetrameric arrangement of the particles. The enlarged class average shows the four protomeric subunits marked using red, blue, green, and magenta-coloured arrows.

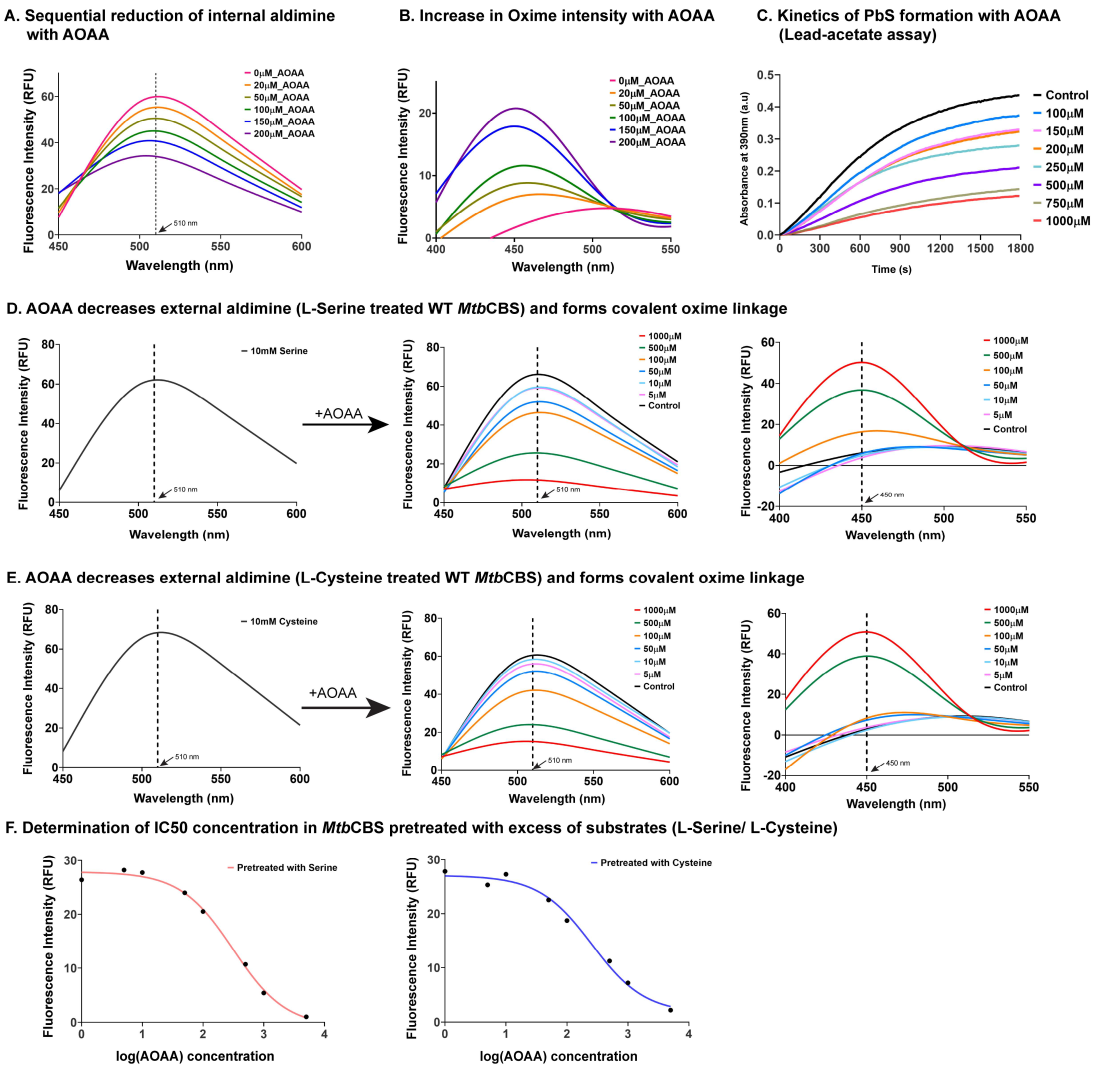


**Fig. S3. Monitoring aldimine, oxime, and enzymatic activity changes along with the dose-dependent effect of AOAA.** Fluorescence emission studies of WT *Mtb*CBS (10μM) with increasing AOAA concentration (20-200μM) show **A.** sequential reduction of aldimine species (($\lambda_{ex}$=410nm) and shift of peak intensity towards **B.** oxime region ($\lambda_{ex}$=350nm). **C.** Inhibition of enzymatic activity with increasing AOAA concentration (100-1000μM) through lead-acetate-based kinetic assay. **D.** Aldimine measurements from *Mtb*CBS incubated with 10mM Serine, followed by the addition of AOAA (5-1000μM) and subsequent measurements of fluorescence emission upon exciting at 410nm (Aldimine) and 350nm (Oxime). The wavelength corresponding to the peak emission intensity is marked by the black dashed vertical line. **E.** Similar fluorescence-based quantification in L-Cysteine pre-treated *Mtb*CBS samples and their emission spectra after AOAA treatment. **F.** The IC50 of AOAA for inhibition of *Mtb*CBS pre-treated with 10mM Serine (red)/ 10mM Cysteine (blue) was obtained by monitoring the aldimine peak intensity changes. Each data point is the average from 2 independent sets.

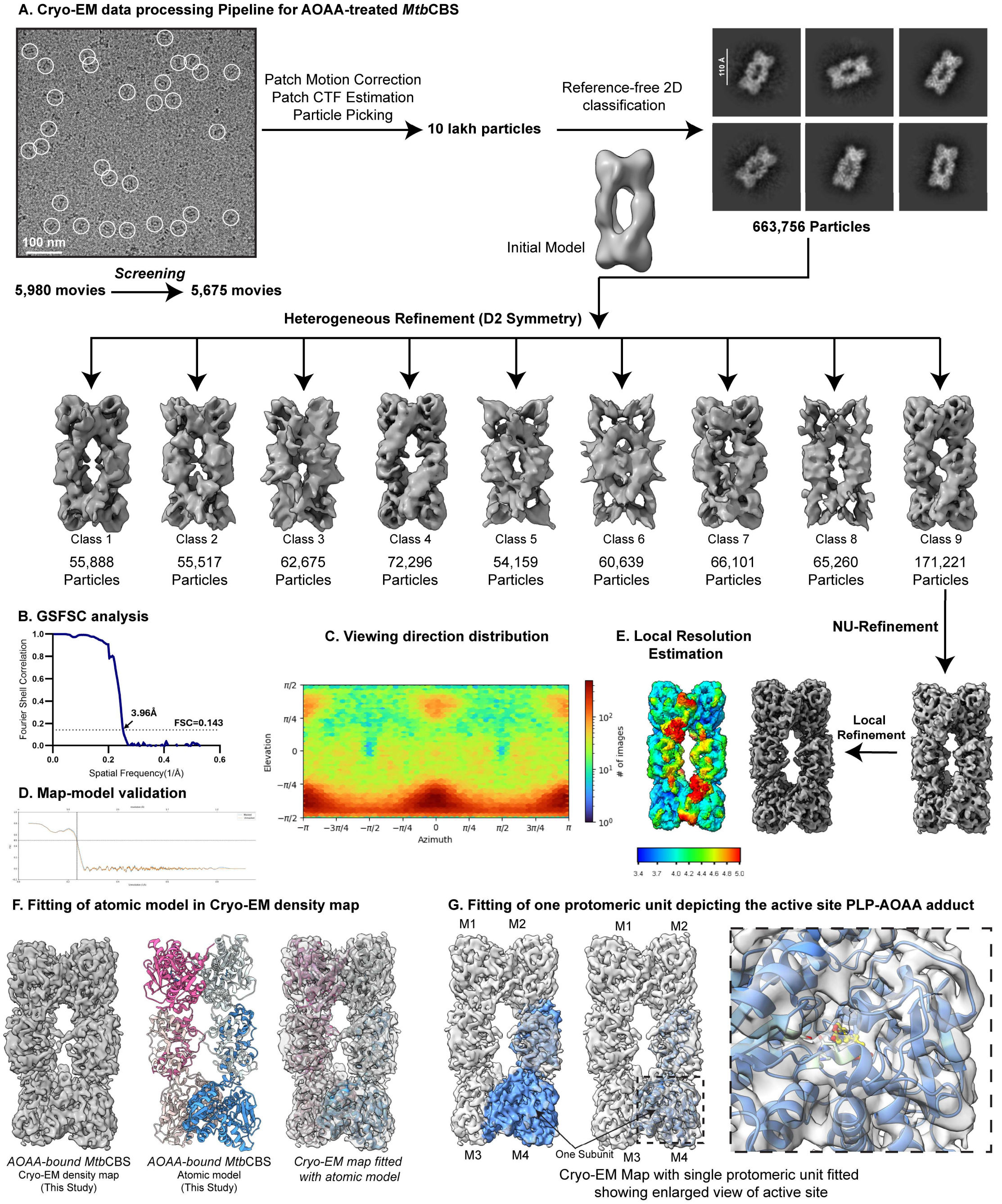


**Fig. S4. Cryo-EM data processing of AOAA-treated *Mtb*CBS. A.** The processing pipeline followed in Cryosparc (as mentioned in Methods) along with the representative cryo-EM micrograph (With particles circled) and reference-free 2D class averages. **B.** GSFSC curve showing the resolution. **C.** Angular distribution plot. **D.** Map-model validation plot. **E.** Cryo-EM density map coloured according to local resolution. **F.** Fitting of the atomic model (PDB ID 9U7O) in cryo-EM density map (EMD-63941). **G.** Fitting of one protomeric subunit depicting the active site PLP-AOAA adduct.

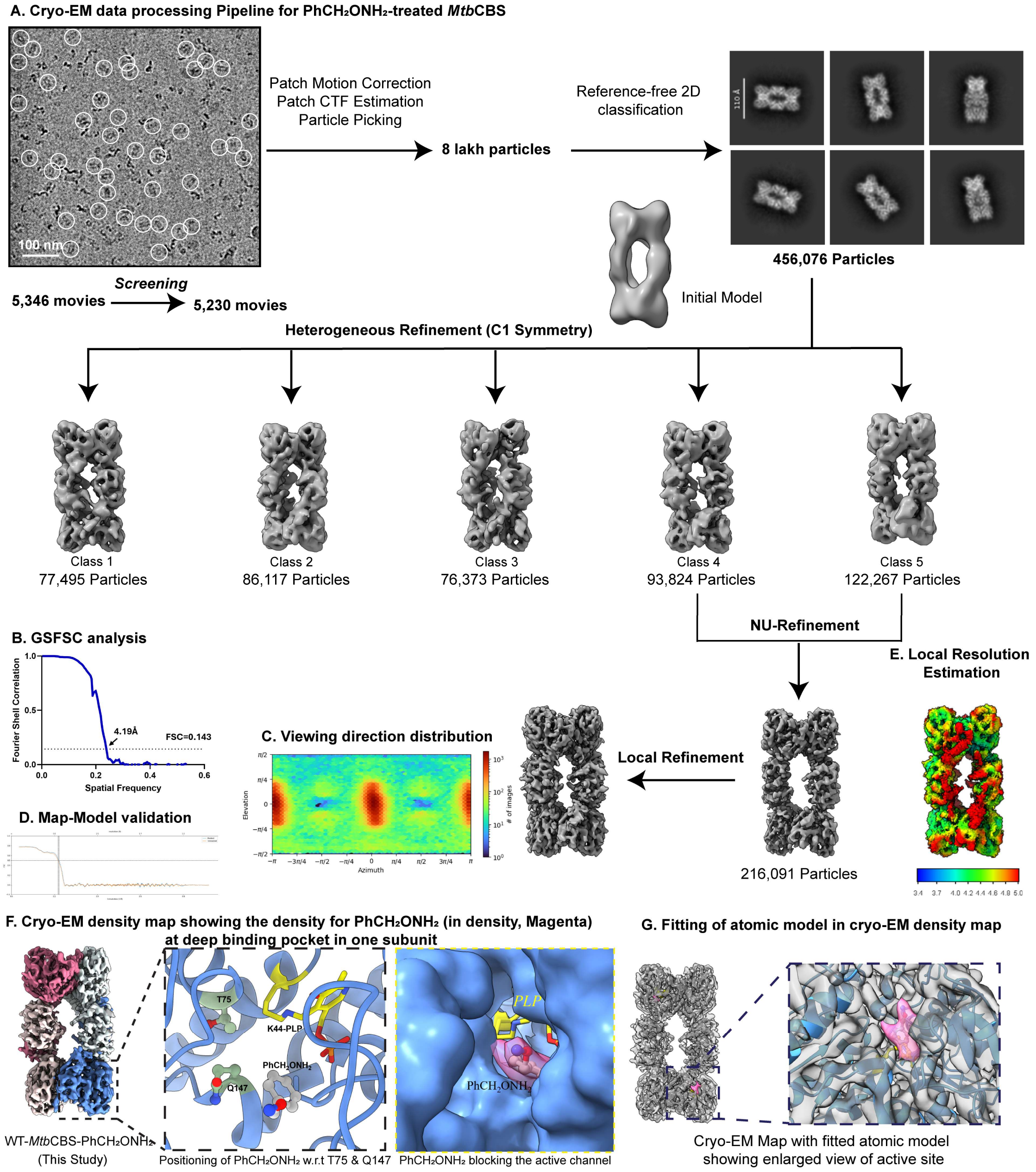


**Fig. S5. Cryo-EM image processing of O-Benzylhydroxylamine-bound tetrameric *Mtb*CBS. A.** Representative cryo-EM micrographs (With particles circled) and calculated reference-free 2D class averages were depicted along with the cryo-EM data processing flow chart. **B.** GSFSC analysis. **C.** Angular distribution plot. **D.** Map-model validation plot. **E.** Cryo-EM density map coloured according to local resolution. **F.** Cryo-EM density map showing the density for $PhCH_2ONH_2$ (in Magenta) at the deep binding pocket in one subunit. **G.** Fitting of the generated atomic model (PDB ID 9U7N) in EMDB map (EMD-63940).

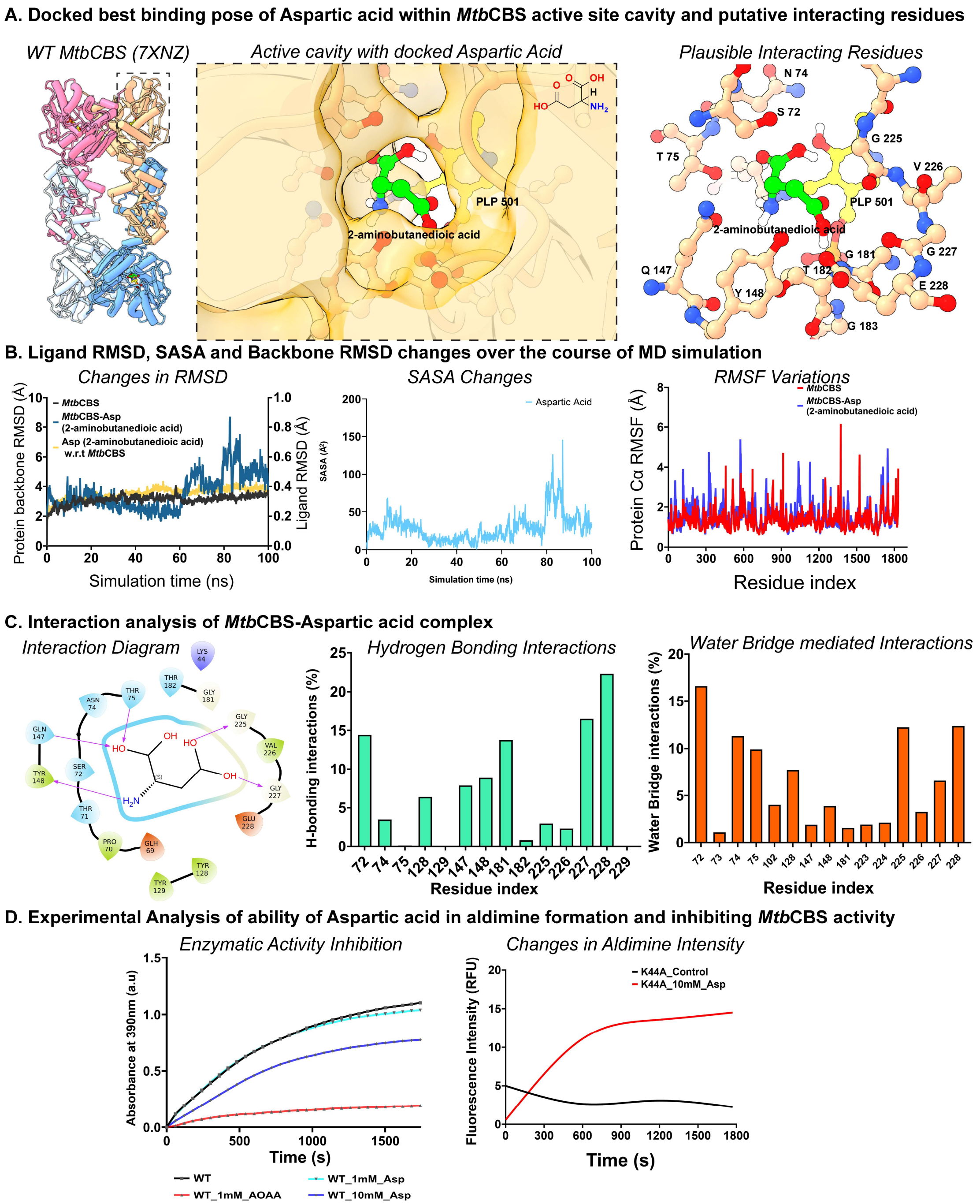


**Fig. S6. In-silico investigations and enzymatic studies using 2-aminobutanedioic acid with *Mtb*CBS. A.** Docked lowest pose of 2-aminobutanedioic acid on PDB structure of tetrameric *Mtb*CBS (7XNZ). The active site region marked with a dashed line is enlarged, showing the obtained lowest energy pose of 2-aminobutanedioic acid enclosed within the surface of *Mtb*CBS. Ball and stick representation showing the plausible interacting residues with the 2-aminobutanedioic acid ligand. **B.** Plots showing the changes in protein backbone RMSD, ligand SASA and protein C-alpha RMSF in the *Mtb*CBS-docked 2-aminobutanedioic acid complex over 100ns simulation time in relation to the uncomplexed control. **C.** The Ligand interaction diagram shows the interaction of 2-aminobutanedioic acid with other polar amino acid residues. The bar plots show the percentage distribution of amino acids in the total number of Hydrogen bonding and water bridge-mediated interactions with the docked ligand over the simulation time. **D.** Lead-acetate-based absorbance ($A_{390nm}$) kinetics with 10μM *Mtb*CBS WT in the presence of 10mM 2-aminobutanedioic acid and 50mM Cysteine showing no significant inhibition in $H_2S$ production and the fluorescence-based kinetic study of 10μM *Mtb*CBS K44A mutant with 10mM 2-aminobutanedioic acid reveals external aldimine formation over time ($\lambda_{ex}$=410nm and $\lambda_{em}$=510nm).

A. Docking and MD-Simulation analysis of O-Benzylhydroxylamine

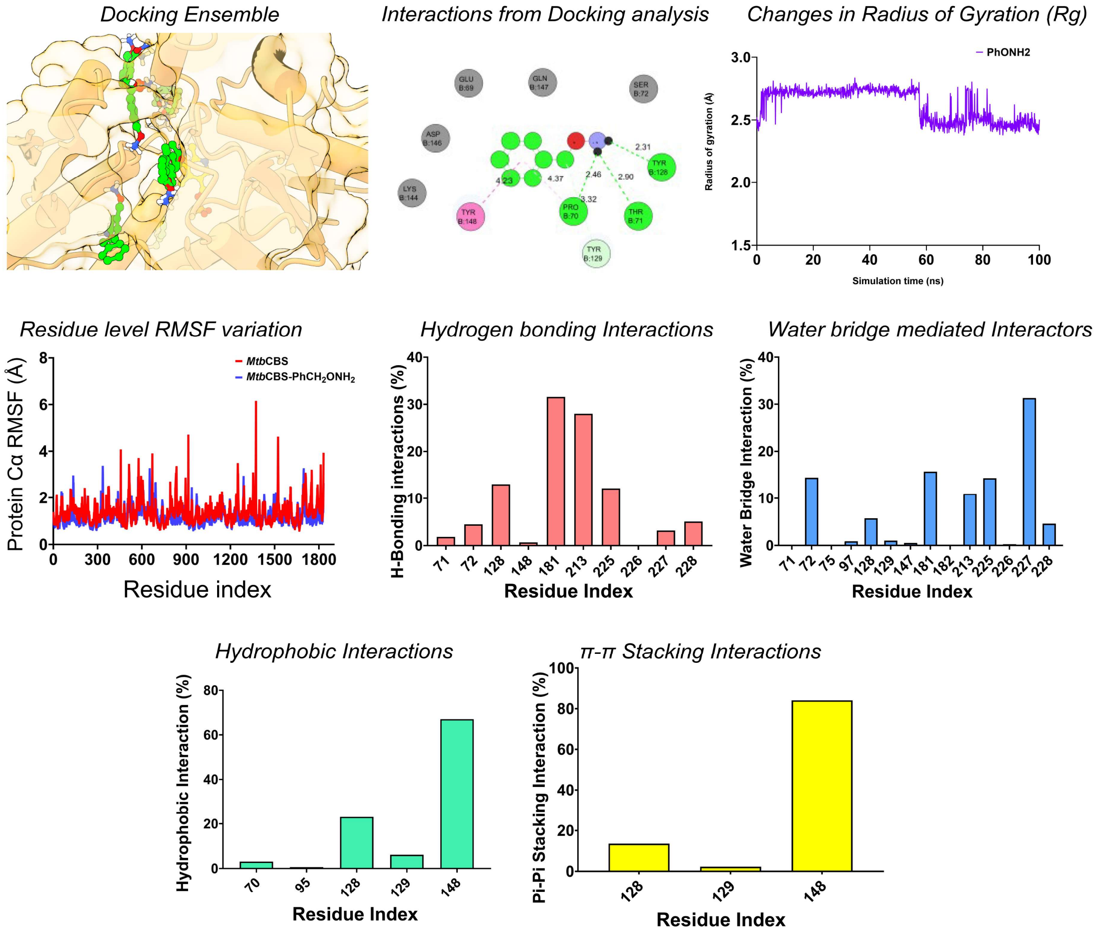


B. Ensemble showing docked conformations of Aspartic Acid along with the modes of interaction and Rg changes

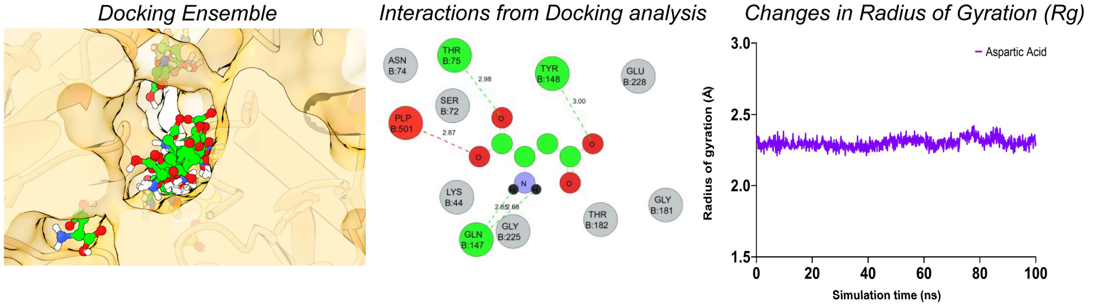


C. Interaction and RMSF changes in O-Benzylhydroxylamine and Aspartic Acid

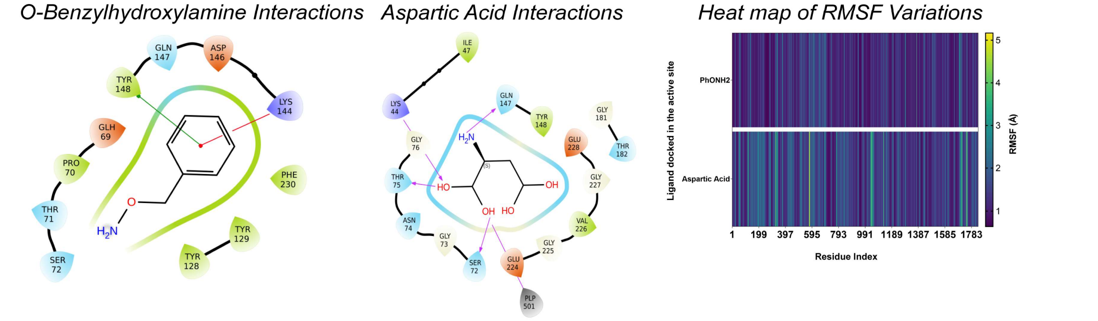

**Fig. S7. Binding of 2-aminobutanedioic acid and O-Benzylhydroxylamine in the active site of *Mtb*CBS (An In-Silico Investigation). A.** Surface representation of *Mtb*CBS active site cavity showing the ensemble of docked poses of 2-aminobutanedioic acid in ball and stick representation. The 2D ligand interaction diagram reveals the interaction of terminal carboxylate with other active site residues (Discovery Studio) and the radius of gyration ($R_g$) changes (average 2.28Å) over the 100ns simulation time. Line plot showing the changes in residue level $C_\alpha$ fluctuations in the presence (blue) and absence (red) of docked $PhCH_2ONH_2$ ligand. Bar plots depict the percentage of residue contribution to hydrogen bonding (brick red), water bridge (cornflower blue), hydrophobic interactions (light green) and π-π stacking interactions (yellow). **B.** Similar representations show the obtained docked conformations of O-Benzylhydroxylamine along with the interaction diagram the extent of ligand extendedness ($R_g$). **C.** Interactions and heatmap showing the residue level $C_\alpha$ fluctuations in simulations of *Mtb*CBS docked with compounds O-Benzylhydroxylamine and 2-aminobutanedioic acid.

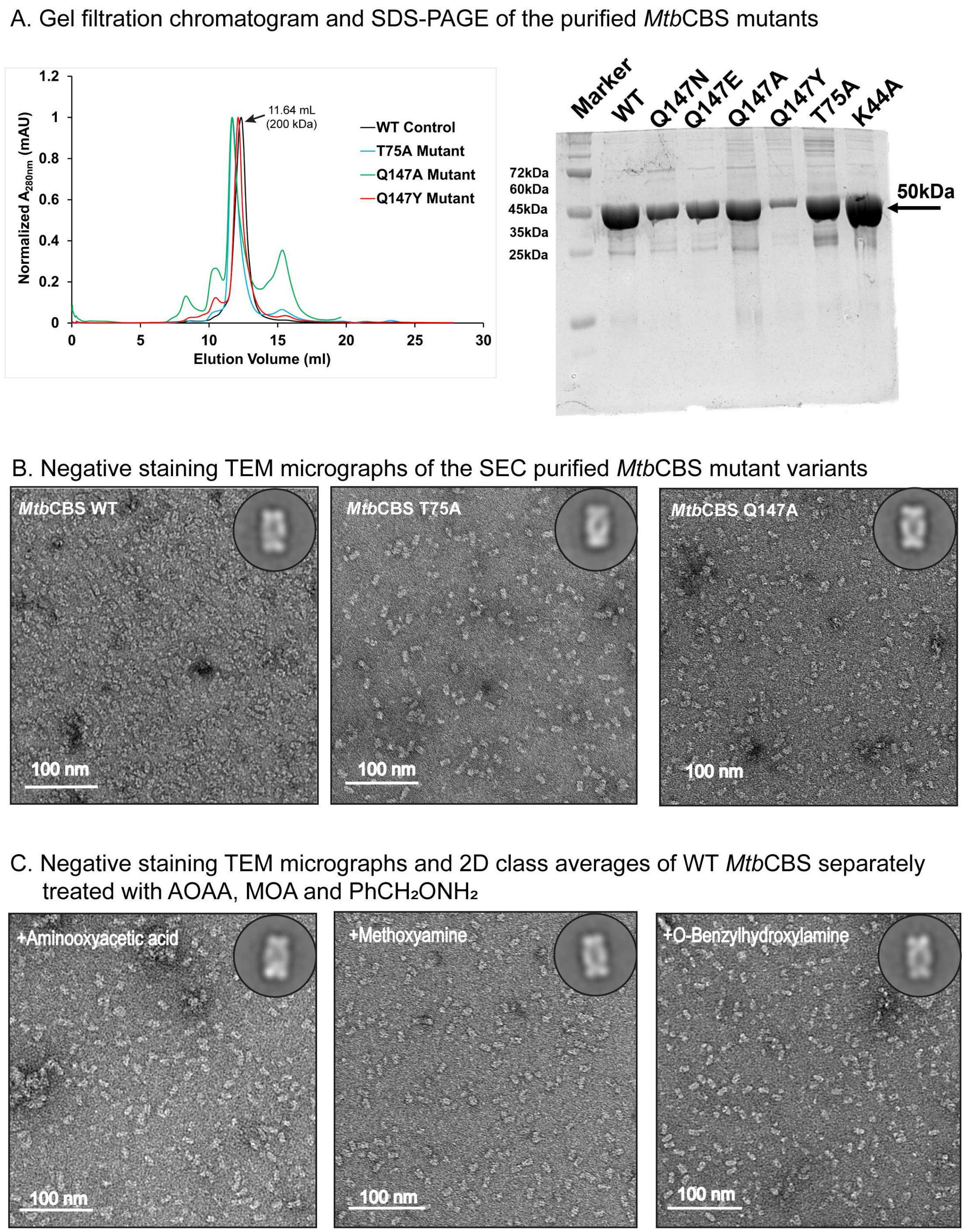


**Fig. S8. Size-exclusion chromatography and NS-TEM studies suggest the tetrameric state of purified recombinant *Mtb*CBS mutant variants and intact WT *Mtb*CBS in the presence of inhibitors. A.** The gel filtration profile of recombinant *Mtb*CBS mutant variants (T75A, Q147A and Q147Y) was obtained using the S200 column. The measured absorbance (A280nm) is normalized (mAU, milli-arbitrary units), and all the mutant variants were eluted at 11.64 ml, corresponding to the tetrameric molecular weight of *Mtb*CBS. 12% SDS-PAGE (in right) shows the protein bands at ~50 kDa corresponding to the molecular mass of *Mtb*CBS mutants. **B.** Representative Negative Staining (NS-TEM) micrographs of the SEC purified mutants of *Mtb*CBS showing isolated tetramers in different orientations (With 2D class averages in top-right corner). **C.** Representative Negative Staining (NS-TEM) micrographs and reference-free 2D class averages of WT *Mtb*CBS treated separately with 1mM AOAA, 1mM MOA and 1mM $PhCH_2ONH_2$ (2D class averages in top-right corner) indicating unaltered overall tetrameric arrangement of *Mtb*CBS.
